# Modeling site-specific amino-acid preferences deepens phylogenetic estimates of viral divergence

**DOI:** 10.1101/302703

**Authors:** Sarah K. Hilton, Jesse D. Bloom

## Abstract

Molecular phylogenetics is often used to estimate the time since the divergence of modern gene sequences. For highly diverged sequences, such phylogenetic techniques sometimes estimate surprisingly recent divergence times. In the case of viruses, independent evidence indicates that the estimates of deep divergence times from molecular phylogenetics are sometimes too recent. This discrepancy is caused in part by inadequate models of purifying selection leading to branch-length underestimation. Here we examine the effect on branch-length estimation of using models that incorporate experimental measurements of purifying selection. We find that models informed by experimentally measured site-specific amino-acid preferences estimate longer deep branches on phylogenies of influenza virus hemagglutinin. This lengthening of branches is due to more realistic stationary states of the models, and is mostly independent of the branch-length-extension from modeling site-to-site variation in amino-acid substitution rate. The branch-length extension from experimentally informed site-specific models is similar to that achieved by other approaches that allow the stationary state to vary across sites. However, the improvements from all of these site-specific but time-homogeneous and site-independent models are limited by the fact that a protein’s amino-acid preferences gradually shift as it evolves. Overall, our work underscores the importance of modeling site-specific amino-acid preferences when estimating deep divergence times—but also shows the inherent limitations of approaches that fail to account for how these preferences shift over time.

## Introduction

Molecular phylogenetics is commonly used to estimate the historical timing of evolutionary events (Yang and Rannala, 2012). This is done by estimating branch lengths based on the inferred number of substitutions, and then converting these branch lengths into units of time under the assumption of a molecular clock (Zuckerkandl and Pauling, 1965; Drummond et al., 2006). However, phylogenetic estimates of the divergence times of many viral lineages are clearly too recent (Duchêne et al., 2014; Ho et al., 2015; Aiewsakun and Katzourakis, 2016). For example, the integration of filoviruses into their host genomes indicate that Ebola and Marburg virus diverged from their common ancestor 7 to 12 million years ago—but the estimate of this divergence time based on phylogenetic analyses of the viral sequences is only ∼ 10,000 years ago (Carroll et al., 2013; Taylor et al., 2014). Similarly, the phylogenetic estimate of when major simian immunodeficiency virus groups diverged is almost 100 times more recent than the estimate based on the geographic isolation of their host species (Wertheim and Worobey, 2009; Worobey et al., 2010). These examples, along with other similar discrepancies with measles virus (Furuse et al., 2010), coronavirus (Wertheim et al., 2013), and hepatitis B virus (Fares and Holmes, 2002; Holmes, 2003), indicate that phylogenetic methods have a systematic bias toward underestimation of deep branches.

This underestimation occurs in part because phylogenetic models do a poor job of describing the real natural selection on protein-coding genes. These genes evolve under purifying selection to maintain the structure and function of the proteins they encode. In general, these constraints are highly idiosyncratic among sites (Echave et al., 2016). However, most phylogenetic models try to account for these constraints using relatively simple approaches such as allowing the rate of substitution to vary across sites according to some statistical distribution (Yang, 1994; Yang et al., 2000). These models of purifying selection are usually inadequate (Duchêne et al., 2015b,a), potentially causing branch lengths to be severely underestimated (Wertheim and Kosakovsky Pond, 2011; Halpern and Bruno, 1998).

More recent work has used mutation-selection models to better account for purifying selection (Halpern and Bruno, 1998; Yang and Nielsen, 2008; Rodrigue et al., 2010; Tamuri et al., 2012; McCandlish and Stoltzfus, 2014). These models explicitly incorporate the fact that different protein sites prefer different amino acids, and so can improve phylogenetic estimates when there are deep branches (Philippe and Laurent, 1998; Lartillot et al., 2007; Le et al., 2008; Quang et al., 2008; Wang et al., 2008). However, these approaches require inferring the site-specific purifying selection from natural sequence data.

Even more recently, it has become possible to directly measure purifying selection on proteins using deep mutational scanning. This high-throughput approach involves experimentally measuring how each amino-acid mutation affects protein function in the lab (Fowler and Fields, 2014). The resulting experimental measurements of which amino acids are preferred at each protein site can be used to inform phylogenetic substitution models (Bloom, 2014a). These experimentally informed codon models (ExpCMs) generally exhibit much better phylogenetic fit than standard substitution models (Doud et al., 2015; Hilton et al., 2017; Haddox et al., 2018; Lee et al., 2018).

Here we examine how ExpCMs and other models of purifying selection estimate branch lengths on a phylogenetic tree of influenza virus hemagglutinin (HA). We find that ExpCMs estimate longer deep branches, and show that this extension of branch length is mostly independent and additive with that achieved by the more conventional approach of modeling rate variation. We also show that ExpCMs estimate similar branch lengths to a mutation-selection model that infers the amino-acid preferences from the natural sequence data rather than using values obtained in experiments. However, all of these mutation-selection models are limited by their failure to account for another feature of purifying selection: the fact that a site’s amino-acid preferences shift over time due to epistasis. Therefore, truly accurate analyses of deep phylogenies need to account for the fact that amino-acid preferences vary across time as well as across sites.

## Results

### Different ways substitution models account for purifying selection

Here we consider how purifying selection is handled by codon models, which are the most sophisticated of the three classes (nucleotide, codon, and amino acid) of phylogenetic substitution models in widespread use for protein-coding genes (Arenas, 2015). Standard codon models distinguish between two types of substitutions: synonymous and nonsynonymous. The relative rate of these substitutions is referred to as dN/dS or *ω*. In their simplest form, codon substitution models fit a single *ω* that represents the gene-wide average fixation rate of nonsynonymous mutations relative to synonymous ones. Here we will use such substitution models in the form proposed by Goldman and Yang (1994). When these models have a single gene-wide *ω* they are classified as M0 by Yang et al. (2000). We will refer to M0 Goldman-Yang models simply as GY94 models (Equation 1). The gene-wide *ω* is usually < 1 (Murrell et al., 2015), and crudely represents the fact that many amino-acid substitutions are under purifying selection.

A single gene-wide *ω* ignores the fact that purifying selection is heterogeneous across sites. The most common strategy to ameliorate this defect is to allow *ω* to vary among sites according to some statistical distribution (Yang, 1994; Yang et al., 2000). For instance, in the M5 variant of the GY94 model (Yang et al., 2000), *ω* follows a gamma distribution as shown in Figure 1A. We will denote this model as GY94+Γ*ω*. A GY94+Γ*ω* captures the fact that the rate of nonsynonymous substitution can vary across sites. However, these models do not capture the fact that the same amino-acid mutation can have very different effects at different sites.

**Figure 1:**
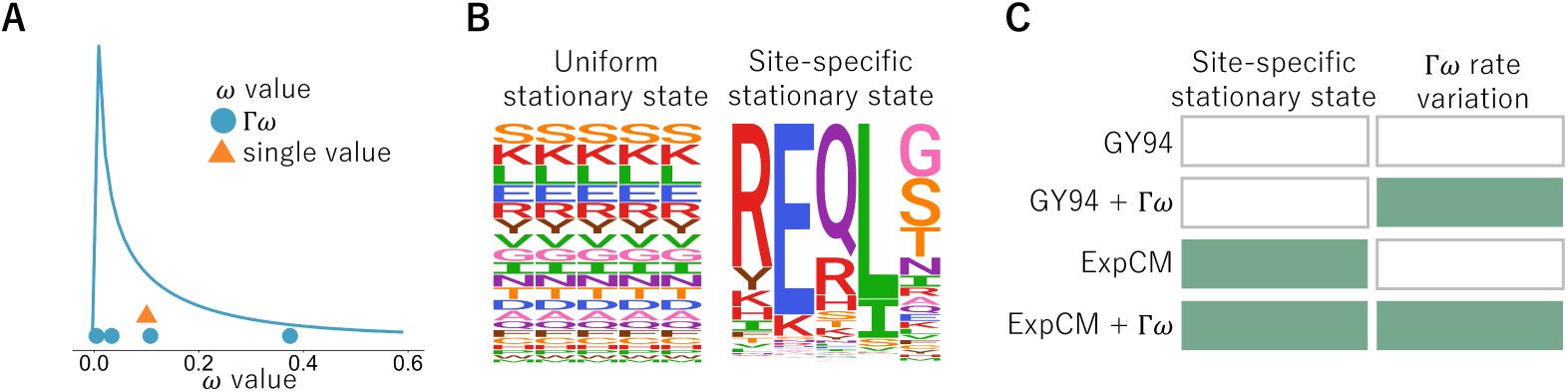
Different ways codon models account for purifying selection. (A) The dN/dS parameter, *ω*, can be defined as one gene-wide average (orange triangle) or allowed to vary according to some statistical distribution (blue line). For computational tractability, the distribution is discretized into *K* bins and *ω* takes on the mean of each bin (blue circles) (Yang, 1994; Yang et al., 2000). A gamma distribution (denoted by Γ) with *K* = 4 bins is shown here. (B) A substitution model’s stationary state defines the expected sequence composition after a very long evolutionary time. Most substitution models have stationary states that are uniform across sites. However, substitution models can have site-specific stationary states. In the logo plots, each column is a site in the protein and the height of each letter is the frequency of that amino acid at stationary state. (C) Substitution models can incorporate neither, one, or both of these features. Here we will use substitution models from the Goldman-Yang (GY94; Goldman and Yang, 1994; Yang et al., 2000) and experimentally informed codon model (ExpCM; Hilton et al., 2017) families with and without gamma-distributed *ω* to represent all possible combinations.

Mutation-selection models account for the fact that purifying selection depends idiosyncratically on the specific amino-acid mutation at each site (Halpern and Bruno, 1998; Yang and Nielsen, 2008; Rodrigue et al., 2010; Tamuri et al., 2012; McCandlish and Stoltzfus, 2014). Here we will consider mutation-selection models where the site-specific selection is assumed to act solely at the protein level (different codons for the same protein are treated as selectively equivalent). Such models explicitly define a different set of amino-acid preferences at each site in the protein. This more mechanistic formulation results in a site-specific stationary state (Figure 1B). These models capture the site-to-site variation in amino-acid composition that is an obvious features of real proteins, and usually better describe actual evolution than models with only rate variation as assessed by Bayesian or maximum-likelihood criteria (Lartillot and Philippe, 2004; Le et al., 2008; Quang et al., 2008; Wang et al., 2008; Rodrigue et al., 2010; Bloom, 2014a,b; Hilton et al., 2017).

However, the increased realism of mutation-selection models comes at the cost of an increased number of parameters. Codon substitution models with uniform stationary states have only a modest number of parameters that must be fit from the phylogenetic data. For instance, a GY94+Γ*ω* model with the commonly used F3X4 stationary state has 12 parameters: two describing the shape of the gamma distribution over *ω*, a transition-transversion rate, and nine parameters describing the nucleotide composition of the stationary state. However, mutation-selection models must additionally specify 19 parameters defining the amino-acid preferences for *each* site (there are 20 amino acids whose preferences are constrained to sum to one). This corresponds to 19 × *L* parameters for a protein of length *L*, or 9,500 parameters for a 500-residue protein. It is challenging to obtain values for these amino-acid preference parameters in a maximum-likelihood framework without overfitting the data (Rodrigue, 2013). Here we will primarily use experimentally informed codon models (ExpCMs), which define the site-specific amino-acid preference parameters *a priori* from deep mutational scanning experiments so that they do not need to be fit from phylogenetic data (see Methods and Bloom, 2014a; Hilton et al., 2017; Bloom, 2017). Because the amino-acid preference parameters in an ExpCM are obtained from experiments, the number of ExpCM free parameters is similar to a non-site-specific substitution model. An alternative strategy of obtaining the amino-acid preference parameters via Bayesian inference (Lartillot and Philippe, 2004; Rodrigue and Lartillot, 2014) is discussed in the last section of the Results.

Importantly, these two strategies for modeling purifying selection are not mutually exclusive. Mutation-selection models such as an ExpCM can still incorporate an *ω* parameter, which now represents the relative rate of nonsynonymous to synonymous substitution *after* accounting for the constraints due to the site-specific amino-acid preferences (Bloom, 2017; Rodrigue and Lartillot, 2017). This *ω* parameter for an ExpCM can be drawn from a statistical distribution (e.g., a gamma distribution) just like for GY94-style models (Rodrigue and Lartillot, 2014; Haddox et al., 2018). We will denote such models as ExpCM+Γ*ω*. Figure 1C shows the full spectrum of models that incorporate all combinations of gamma-distributed *ω* and site-specific stationary states.

### Effect of stationary state and rate variation on branch-length estimation

Given a single branch, a substitution model transforms sequence identity into branch length. Under a molecular-clock assumption, this branch length is proportional to time. The transformation from sequence identity to branch length is trivial when the sequence identity is high. For instance, when there has only been one substitution, then the sequence identity will simply be
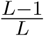
for a gene of *L* sites, and even a simple exponential model (Zuckerkandl and Pauling, 1965) will correctly infer the short branch length of 1/*L* substitutions per site. However, as substitutions accumulate it becomes progressively more likely for multiple changes to occur at the same site. In this regime, the accuracy of the substitution model becomes critical for transforming sequence identity into branch length. Any time-homogenous substitution model predicts that after a very large number of substitutions, two related sequences will approach some asymptotic amino-acid sequence identity. For instance, if all 20 amino acids are equally likely in the stationary state, then this asymptotic sequence identity will be
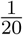
=0.05. If the substitution model underestimates the asymptotic sequence identity then it will also underestimate long branch lengths, since it will predict that sequences that have evolved for a very long time should be more diverged than is actually the case.

Figure 2 shows how different substitution models predict amino-acid sequence identity to decrease as a function of branch length using model parameters fit to a phylogeny of H1 influenza hemagglutinin (HA) genes. The GY94 model predicts the same behavior for all sites, since it does not have any site-specific parameters, with an asymptotic sequence identity of 0.062. While this predicted sequence identity is higher than
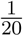
= 0.05 due to redundant codon and nucleotide biases favoring certain amino acids, it is much lower than the pairwise identity of even the most diverged HAs in nature. While it is of course possible that the identity of HAs in nature would become even lower given more time, it seems biochemically improbable that it would ever become as low as 0.062. The reason is that like many proteins HA has a highly conserved structure and function that imposes constraints that cause many sites to sample only a small subset of the 20 amino acids among all known HA homologs (Nobusawa et al., 1991).

**Figure 2:**
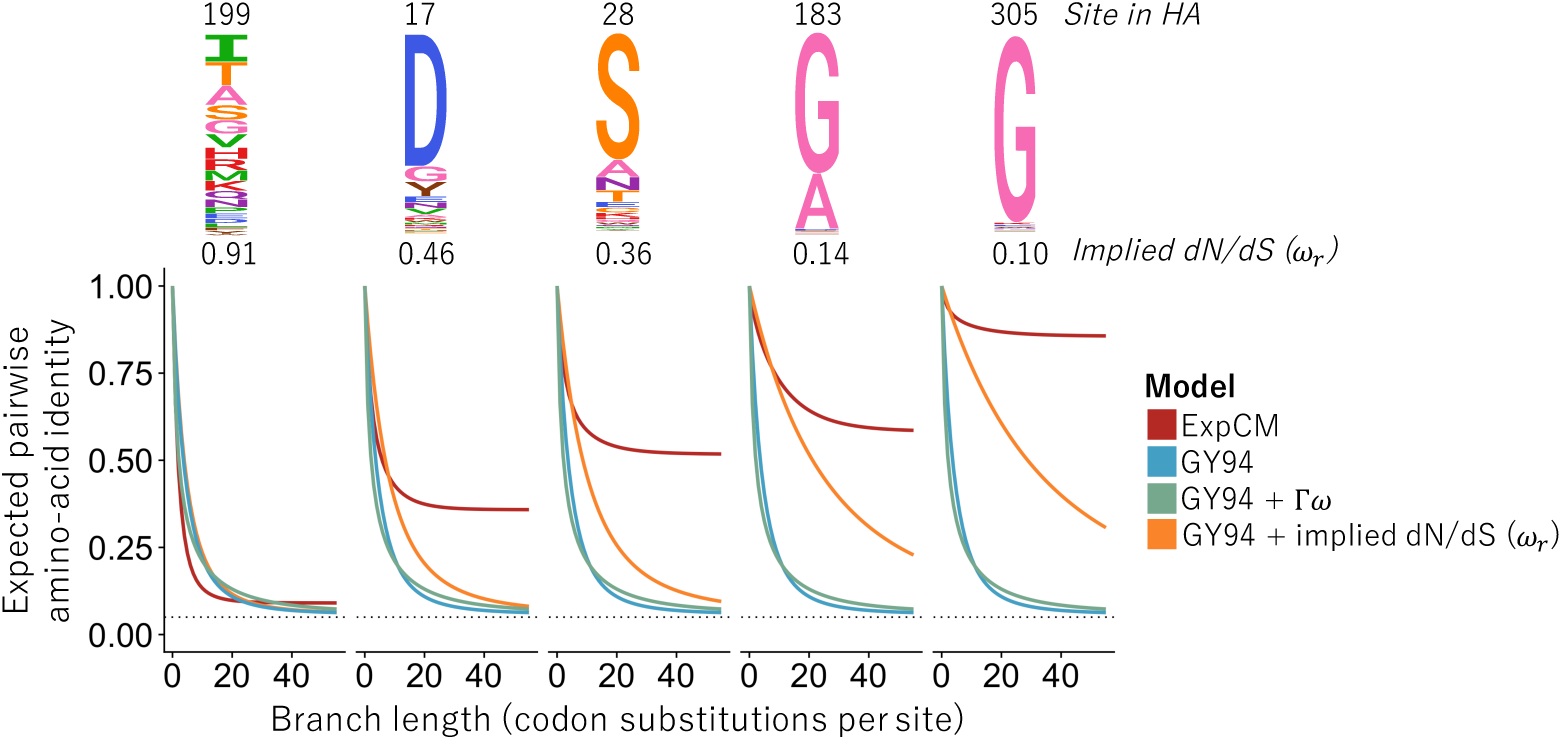
Effect of stationary state and Γ*ω* rate variation on predicted asymptotic sequence divergence. The logo plots at top show the amino-acid preferences for some sites in an H1 influenza hemagglutinin protein as experimentally measured by Doud and Bloom (2016). The graphs show the expected amino-acid identity at that site for two sequences separated by a branch of the indicated length (Equation 9). For the GY94 model, the graphs are identical for all sites since this model does not have site-specific parameters; the same is true for GY94+Γ*ω*. The graphs do differ among sites if we calculate a different *ω_r_* for each site *r* in the GY94 model using the amino-acid preferences (Equation 7; Spielman and Wilke, 2015b). However, all GY94 models, including the one with site-specific *ω_r_* values, approach the same asymptote since they all have the same stationary state. The ExpCM has different asymptotes for different sites since it accounts for how amino-acid preferences lead to site-specific stationary states.

Accounting for site-to-site dN/dS rate variation in GY94 models affects the rate at which the asymptotic sequence identity is approached, but not the actual value of this asymptote. For instance, Figure 2 shows that the GY94+Γ*ω* model takes longer to reach the asymptote than GY94, but that the asymptote is identical for both models. This fact holds true even if we use experimental measurements of HA’s site-specific amino-acid preferences (Doud and Bloom, 2016) to calculate a different *ω_r_* value for each site using the method of Spielman and Wilke (2015b) (see Equation 7). Specifically, this GY94+*ω_r_* model predicts that different sites will approach the asymptote at different rates, but the asymptote is always the same (Figure 2). The invariance of the asymptotic sequence identity under different schemes for modeling *ω* is a fundamental feature of the mathematics of reversible substitution models. These models are reversible stochastic matrices, which can be decomposed into stationary states and symmetric exchangeability matrices (Nielsen, 2006). The stationary state is invariant with respect to multiplication of the symmetric exchangeability matrix by any non-zero number. Different schemes for modeling *ω* only multiply elements of the symmetric exchangeability matrix. Therefore, no matter how “well” a model accounts for site-to-site variation in *ω*, it will always have the same stationary state as a simple GY94 model.

However, mutation-selection models such as ExpCMs have site-specific stationary states. They predict that different sites will have different asymptotic sequence identities (Figure 2)—a prediction that accords with the empirical observation that some sites are much more variable than others in alignments of highly diverged sequences. For instance, Figure 2 shows that at sites such as 183 and 305 in the H1 HA, an ExpCM but not a GY94-style model predicts that the identity will always be relatively high. When sites with highly constrained amino-acid preferences such as these are common, an ExpCM can estimate a long branch length at modest sequence identities that a GY94 model might attribute to a shorter branch.

### Simulations demonstrate how failure to model site-specific amino-acid preferences leads to branch-length underestimation

To directly demonstrate the effect of stationary state and Γ*ω* rate variation on branch-length estimation, we tested the ability of a variety of models to accurately infer branch lengths on simulated data (Figure 3). Specifically, we simulated alignments of sequences along the HA phylogenetic tree using an ExpCM parameterized by the amino-acid preferences of H1 HA as experimentally measured by deep mutational scanning (Doud and Bloom, 2016). We then estimated the branch lengths from the simulated sequences using all the substitution models in Figure 1C, and compared these estimates to the actual branch lengths used in the simulations. Note that these simulations closely parallel those performed by Halpern and Bruno (1998) and Wertheim and Kosakovsky Pond (2011).

**Figure 3:**
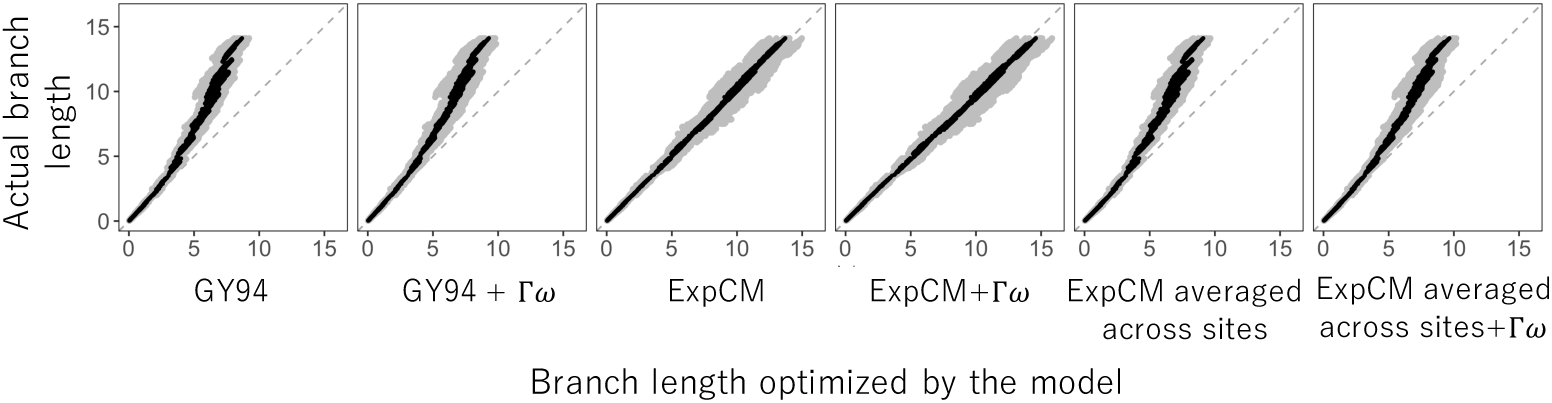
Branch lengths inferred on data simulated under a model with site-specific amino-acid preferences. We simulated alignments along a HA phylogenetic tree using an ExpCM parameterized by the actual site-specific amino-acid preferences for an H1 HA (Doud and Bloom, 2016). We then inferred the branch lengths of this tree from the simulated alignments. The inferred branch lengths for various models are plotted on the x-axis, and the actual branch lengths used in the simulations are on the y-axis. We performed 10 simulations and inferences, and gray points show each inferred branch length from each simulation, and black points show the average of each branch length across simulations. The grey dashed line at *y* = *x* represents what would be seen if the inferred branch lengths exactly matched those used in the simulations.

The models with a uniform stationary state underestimated the lengths of long branches on the phylogenetic tree of the simulated sequences (Figure 3). The GY94 model estimated branch lengths that are ∼60% of the true values for the longest branches. Accounting for site-to-site variation in *ω* did not fix the fundamental problem: the GY94+Γ*ω* did slightly better, but still substantially underestimated the longest branches. However, there was no systematic underestimation of long branches by the ExpCM and ExpCM+Γ*ω* models. The improved performance of the ExpCMs is due to their modeling of the site-specific amino-acid preferences: if we parameterize ExpCMs by amino-acid preferences that have been averaged across HA sites (and so are no longer site-specific), then they perform no better than GY94 models (Figure 3). Therefore, models with uniform stationary states underestimate the length of long branches in phylogenies of sequences that have evolved under strong site-specific amino-acid preferences.

### Experimentally informed site-specific models estimate longer branches on real data

The foregoing section shows the superiority of ExpCMs to GY94 models for estimating long branches on phylogenies simulated with ExpCMs. But how do these models perform on real data? Real genes do evolve under functional constraint, but these constraints are almost certainly more complex than what is modeled by an ExpCM. However, if ExpCMs do a substantially better job than GY94 models of capturing the true constraints, then we might still expect them to estimate more accurate branch lengths.

To test the models on real data, we used actual sequences of influenza HA. The topology of HA phylogenetic trees makes these sequences an interesting test case for branch-length estimation. HA consists of a number of different subtypes. Sequences within a subtype have >68% amino-acid identity, but sequences in different subtypes have as little as 38% identity. However, HA proteins from all subtypes have a highly conserved structure that performs a highly conserved function (Ha et al., 2002; Russell et al., 2004). We used RAxML (Stamatakis, 2006) with a nucleotide substitution model (GTRCAT) to infer a phylogenetic tree for 92 HA sequences drawn from 15 of the 18 subtypes (we excluded bat influenza and one other rare subtype). For the rest of this paper, we fix the tree topology to this `RAxML`-inferred tree. Although the nucleotide model used with `RAxML` to infer this tree topology is probably less accurate than codon models, the modular subtype structure of the HA phylogeny means that most of the phylogenetic uncertainty lies in the length of the long branches separating the subtypes rather than in the tree topology itself.

Deep mutational scanning has been used to measure the amino-acid preferences of all sites in two different HAs. One scan measured the preferences of an H1 HA (Doud and Bloom, 2016) and the other measured the preferences of an H3 HA (Lee et al., 2018). The amino-acid preferences measured for these two HAs are shown in Supplementary figure 1 and Supplementary figure 2. The H1 and H3 HAs have only ∼42% amino-acid identity. As described in Lee et al. (2018), the amino-acid preferences clearly differ between the H1 and H3 HA at a substantial number of sites (these differences are apparent in a simple visual comparison of Supplementary figure 1 and Supplementary figure 2; see site 33 as an example). Therefore, we also created a third set of amino-acid preferences by averaging the measurements for the H1 and H3 HAs, under the conjecture that these averaged preferences might better describe the “average” constraint on sites across the full HA tree (Supplementary figure 3). These three sets of HA amino-acid preferences define three different ExpCMs.

We fit the GY94 model and each of the three ExpCMs to the fixed HA tree topology estimated using `RAxML`, and also tested a version of each model with Γ*ω* rate variation. Table 1 shows that all ExpCMs fit the actual data much better than the GY94 models. The best fit was for the ExpCM informed by the average of the H1 and H3 deep mutational scans. For all models, incorporating Γ*ω* rate variation improved the fit, although even ExpCMs without Γ*ω* greatly outperformed the GY94+Γ*ω* model (Table 1). As mentioned in the previous section, *ω* is generally < 1 when a single value is fit to all sites in a gene (Murrell et al., 2015), and this is the case for all the models we tested (Table 1). However, the ExpCMs always fit an *ω* greater than the GY94 model, suggesting that the site-specific amino-acid preferences capture some of the purifying selection that the GY94 models can represent only via a small *ω*. Among the models with Γ*ω*, the GY94+Γ*ω* model fits all four *ω* categories to values ≪ 1, but the ExpCM+Γ*ω* models fit one of the *ω* categories to a value > 1. This increase in *ω* values makes sense given the different interpretation of *ω* for each family of models. The ExpCM *ω* is the relative rate of fixation of nonsynonymous to synonymous mutations *after* accounting for the functional constraints described by the amino-acid preferences. This more realistic null model gives ExpCMs enhanced power to detect diversifying selection for amino-acid change (Bloom, 2017; Rodrigue and Lartillot, 2017), which is known to occur at some sites in HA due to immune selection (Bedford et al., 2014).

**Table 1:** Fitting of substitution models to the HA phylogenetic tree. All ExpCMs describe the evolution of HA better than the GY94 models, as evaluated by the Akaike information criteria (ΔAIC, Posada and Buckley, 2004). The models fit here are the same ones in Figure 4. The *ω* value for each of the *K* = 4 bins is shown for the models with Γ*ω* rate variation. All ExpCMs fit a stringency parameter > 1.

| Model | $\Delta\text{AIC}$ | Log Likelihood | $\omega$ | Stringency parameter ($\beta$) |
| --- | --- | --- | --- | --- |
| ExpCM (H1+H3 avg) + $\Gamma\omega$ | 0 | -51083 | 0.19, 0.50, 0.91, 1.86 | 1.69 |
| ExpCM (H1+H3 avg) | 1063 | -51616 | 0.14 | 1.77 |
| ExpCM (H1) + $\Gamma\omega$ | 1321 | -51744 | 0.12, 0.42, 0.89, 2.13 | 1.11 |
| ExpCM (H3) + $\Gamma\omega$ | 1777 | -51972 | 0.10, 0.36, 0.76, 1.84 | 1.28 |
| ExpCM (H1) | 2670 | -52419 | 0.12 | 1.21 |
| ExpCM (H3) | 3377 | -52773 | 0.12 | 1.43 |
| GY94 + $\Gamma\omega$ | 4817 | -53487 | 0.00, 0.03, 0.08, 0.24 | - |
| GY94 | 7892 | -55025 | 0.07 | - |

Importantly, models that account for purifying selection via either Γ*ω* rate variation or site-specific amino-acid preferences do not just exhibit better fit—they also estimate longer branches on the HA tree. Figure 4 shows the branch lengths optimized by each model on a common scale. The tree’s deepest branches are shortest when they are optimized by the GY94 model, which lacks both Γ*ω* and site-specific amino-acid preferences. Adding either Γ*ω* rate variation or site-specific amino-acid preferences increases the length of the deep branches. Specifically, the tree’s diameter (the distance between the two most divergent tips) for the GY94+Γ*ω* model is 159% of the GY94 model tree diameter (Supplementary table 1). The tree diameter is 122% and 135% of the GY94 model tree diameter for ExpCMs informed by H1 or H3 amino-acid preferences, respectively, and 160% of the GY94 model for the ExpCM informed by the average of the H1 and H3 preferences (Supplementary table 1).

**Figure 4:**
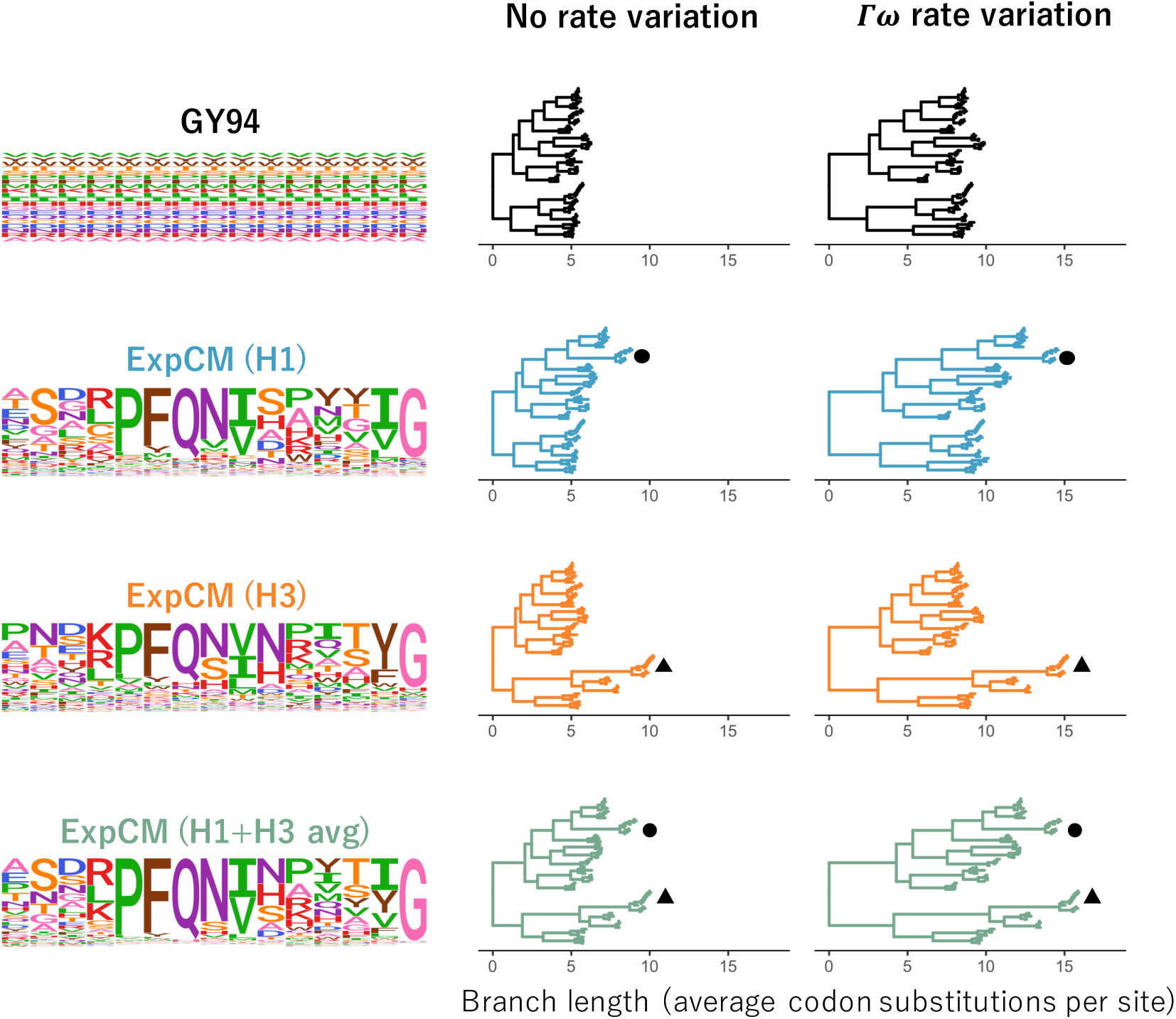
Effect of site-specific amino-acid preferences and Γ*ω* rate variation on HA branch length estimation. The branch lengths of the HA tree were optimized using the indicated ExpCM or GY94 model. The amino-acid preferences defining the model (ExpCM) or implied by the model (GY94) are shown as logo plots for 15 sites in HA; the full set of experimentally measured amino-acid preferences are in Supplementary figure 1, Supplementary figure 2, and Supplementary figure 3. The ExpCMs use amino-acid preferences measured in deep mutational scanning of an H1 HA (Doud and Bloom, 2016), an H3 HA (Lee et al., 2018), or the average of the measurements for these two HAs. Circle denotes the H1 clade and triangle denotes the H3 clade. The root of each tree is placed where it would fall if the tree was midpoint rooted using the branch lengths inferred by `RAxML` using the GTRCAT model. This figure enables qualitative visualization of the trees; for a quantitative comparison of branch lengths optimized by different models, see Figure 5.

The deepening of branch lengths that results from the Γ*ω* and site-specific amino-acid preference approaches to modeling purifying selection are largely independent. This can be seen by examining the ExpCM+Γ*ω* models, which combine Γ*ω* rate variation with site-specific amino-acid preferences. As shown in Figure 4, these ExpCM+Γ*ω* models estimate longer branches than models with just Γ*ω* rate variation (GY94+Γ*ω*) or just site-specific amino-acid preferences (ExpCMs). The near independence of these effects is quantified in Supplementary table 1, which shows that 76% of the tree diameter extension of ExpCM(H1+H3 avg)+Γ*ω* versus can be explained by simply adding the extension from incorporating Γ*ω* (GY94+Γ*ω* versus GY94) to the extension from incorporating site-specific amino-acid preferences (compare ExpCM(H1+H3 avg) to GY94).

However, while adding Γ*ω* rate variation increases the length of deep branches in a roughly uniform fashion across the tree, the branch lengthening from adding site-specific amino-acid preferences is not uniform across the tree (Figure 4 and Figure 5). Instead, the increase in branch length is most pronounced on branches leading to the HA sequence that was used in the deep mutational scanning experiment that informed the ExpCM. For instance, the ExpCM informed by the H1 data most dramatically lengthens branches near the H1 clade of the tree, while the ExpCM informed by the H3 data has the largest effect on branches near the H3 clade. The ExpCM informed by the average of the H1 and H3 data has a more uniform effect across the tree, but still most strongly extends branches leading to either the H1 or H3 clade. Therefore, Figure 4 and Figure 5 show that ExpCMs estimate longer branches, but that the effect is shaped by the set of amino-acid preferences used to inform the model.

**Figure 5:**
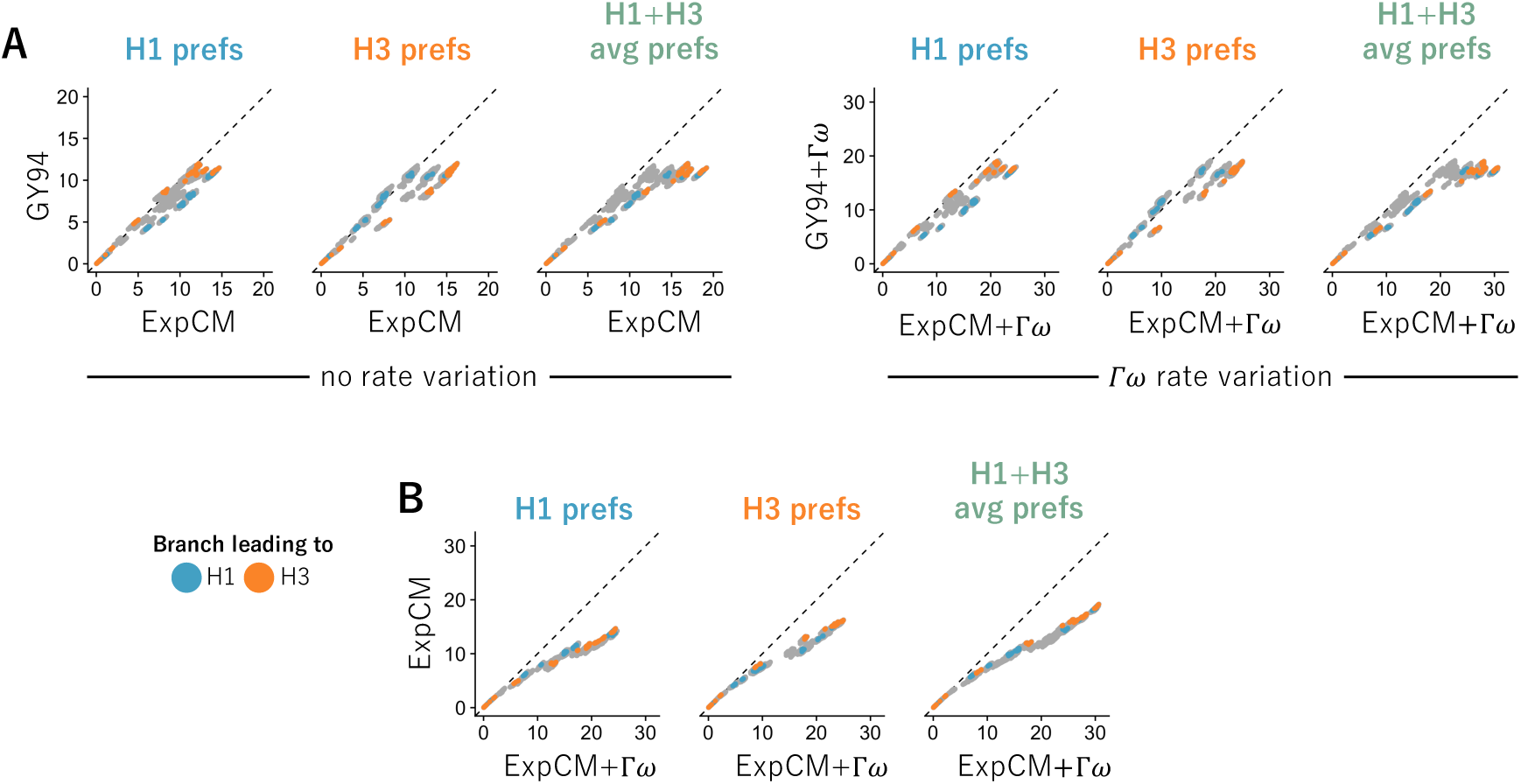
Modeling site-specific amino-acid preferences using deep mutational scanning experiments extends branch lengths, especially for branches leading to the HA used in the experiment. The points indicate the total length of branches separating all pairs of tips on the HA phylogenetic tree when the tree is optimized under the indicated model. Blue and orange denote branches that lead to the H1 and H3 HAs used in the deep mutational scanning. The amino-acid preference set defining the ExpCM is labeled above each each plot. (A) ExpCMs defined by amino-acid preferences from any of the deep mutational scanning experiments estimate generally longer branches than the GY94 model, with the increase particularly pronounced for branches leading to the HA used in the experiment. (B) The addition of Γ*ω* rate variation further extends branch lengths, without any apparent bias towards the HAs used in the experiment. Note that this figure shows the same data as Figure 4 in a different form.

### Shifting amino-acid preferences limit the benefits of models with site-specific stationary states for estimating long branch lengths

The fact that an ExpCM leads to the most profound increase in branch length leading to the sequence used in the experiment can be rationalized in terms of existing knowledge about epistasis during protein evolution. Each ExpCM is informed by a single set of experimentally measured amino-acid preferences. But in reality, the effect of a mutation at one site in a protein can depend on the amino-acid identities of other sites in the protein (Ortlund et al., 2007; Gong et al., 2013; Harms and Thornton, 2014; Tufts et al., 2014; Starr et al., 2018). This epistasis can lead to shifts in a protein’s amino-acid preferences over evolutionary time (Pollock et al., 2012; Doud et al., 2015; Shah et al., 2015; Bazykin, 2015; Haddox et al., 2018). Because the deep mutational scanning experiments that inform our ExpCMs were each performed in the context of a single HA genetic background, their measurements do not account for the accumulation of epistatic shifts in amino-acid preferences as HA evolves. Therefore, an ExpCM is expected to most accurately describe the evolution of sequences closely related to the one used in the experiment.

We can observe how shifting amino-acid preferences degrade the accuracy of an ExpCM by fitting the model to trees containing increasingly diverged sequences. For both H1 and H3 HAs, we created three phylogenetic trees (Supplementary figure 5): a “low” divergence tree that contains sequences with ≥59% amino-acid identity to the HA used in the experiment, an “intermediate” divergence tree that contains sequences with ≥46% amino-acid identity to the HA in the experiment, and a “high” divergence tree that contains all HAs (which have as little as 38% identity to the HA in the experiment). Figure 6 shows the subtrees containing each of these sets of HA sequences. For each subtree, we examined the congruence between site-specific natural selection and the amino-acid preferences measured in the deep mutational scanning experiment using the ExpCM stringency parameter *β* (Bloom, 2014b; Hilton et al., 2017). Values of *β* that are >1 indicate that natural selection prefers the same amino acids as the experiments but with a greater stringency, suggesting strong congruence between natural selection and the experimental preferences. In contrast, values of *β* that are <1 flatten the preferences, suggesting that they provide a relatively poor description of natural selection on the protein.

**Figure 6:**
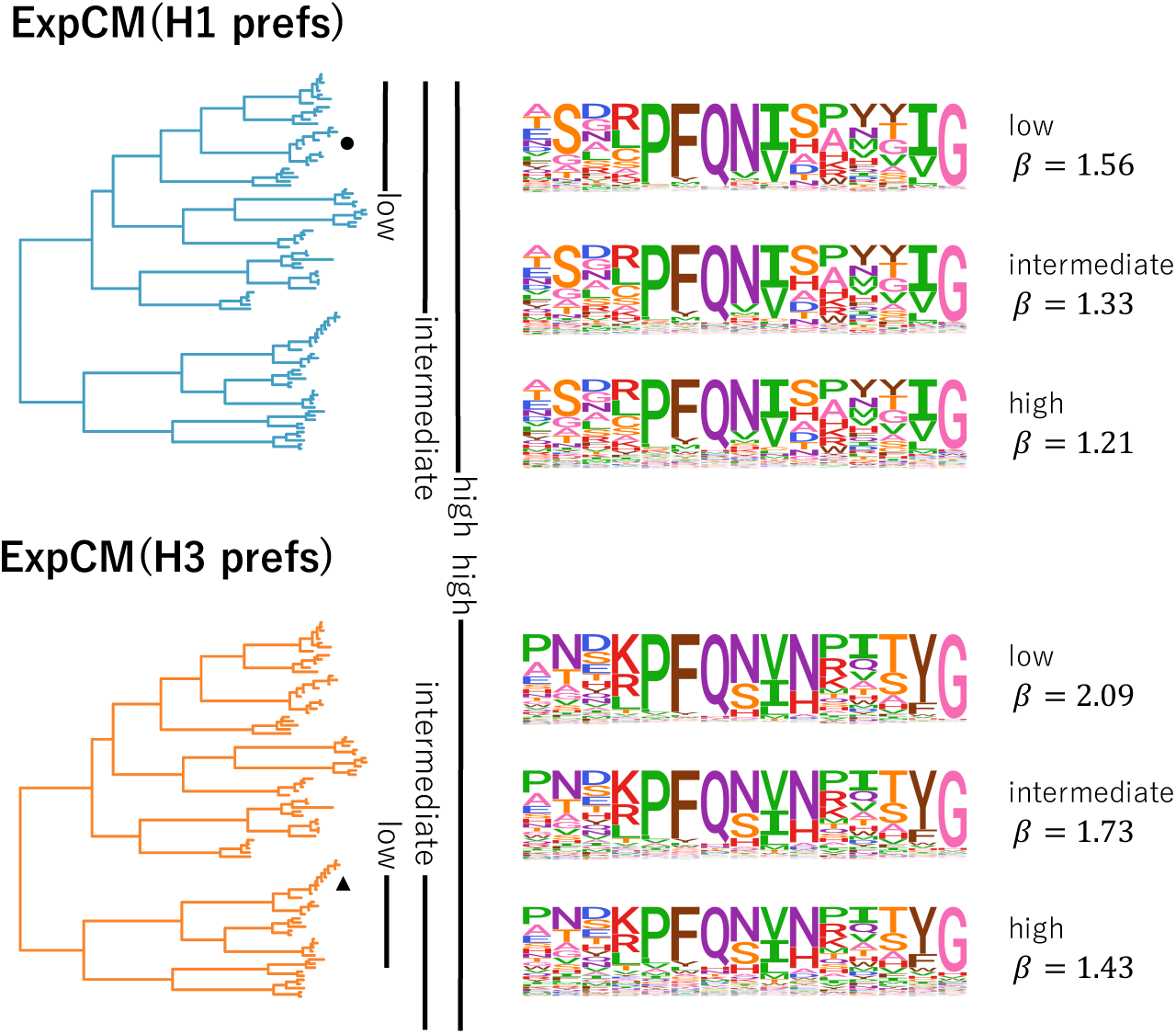
The congruence between natural selection and the deep mutational scanning measurements decreases with sequence divergence. We fit an ExpCM informed by the H1 or H3 deep mutational scanning experiments to trees spanning sequences with low, intermediate, and high divergence from the sequence used in the experiment. The ExpCM stringency parameter (*β*) is a measure of the congruence between natural selection and the experimental measurements (Bloom, 2014b; Hilton et al., 2017). Larger values of *β* indicate that natural selection prefers the same amino acids as the experiments but with greater stringency. As divergence increases between the HA used in the experiment and the other sequences in the tree, the *β* value decreases and the amino-acid preference “flatten.” Therefore, the preferences measured in each experiment are progressively less congruent with natural selection as we include increasingly diverged sequences.

The value of *β* decreases as the divergence from the sequence used in the deep mutational scan increases Figure 6. This inverse relationship between *β* and overall divergence is seen for the ExpCMs informed by both the H1 and H3 experiments. As *β* decreases, the preferences “flatten” and so the ExpCM draws less information from the experiment. At the most extreme value of *β* = 0, the preferences would be perfectly uniform and look similar to the GY94 preferences in Figure 4. In reality, *β* never reaches a value this low, indicating the deep mutational scanning experiments remain somewhat informative about real natural selection across the entire swath of HAs. However, Figure 6 shows that the amino-acid preferences clearly become less informative about natural selection as we move away from the experimental sequence on the tree. This shifting of amino-acid preferences helps explain why the ExpCM informed by the average of the H1 and H3 experiments performs best (Table 1, Figure 4, and Figure 5): averaging the measurements across these two HAs is a heuristic method of accounting for shifts in preferences during HA evolution.

The fact that amino-acid preferences shift as a protein evolves leaves us with an inherent tension: models with site-specific amino-acid preferences only become important for accurate branch-length estimation as sequences become increasingly diverged, but this same divergence degrades the accuracy of extrapolating the amino-acid preferences from any given experiment across the phylogenetic tree. Crucially, this problem is more fundamental than the inability of a single deep mutational scanning experiment to measure amino-acid preferences in more than one genetic background. If amino-acid preferences shift during evolution, there simply will not be any single set of time-homogeneous site-specific preferences that accurately describes evolution along the entirety of a phylogenetic tree that covers a wide span of sequences.

### A model with amino-acid preferences estimated from natural sequences gives similar results to an ExpCM

The previous sections used ExpCMs, which are mutation-selection models that use site-specific amino-acid preferences that have been measured by experiments. However, there are other mathematically similar implementations of mutation-selection models that infer the amino-acid preferences directly from the natural sequence data. When these models are designed for use in phylogenetic inference, they are generally implemented in a Bayesian framework, which avoids the overfitting problems associated with trying to make maximum-likelihood estimates of the thousands of amino-acid preference parameters (Lartillot, 2014). (Note that the maximum-likelihood implementations of Tamuri et al. (2012, 2014) are designed for estimating the amino-acid preferences, not for phylogenetic inference.) The model most comparable to our ExpCMs is the codon mutation-selection model implemented in `PhyloBayes–MPI`, which we will refer to as pbMutSel (Rodrigue and Lartillot, 2014). In the pbMutSel model, the amino-acid preferences are modeled using Dirichlet processes rather than derived from experiments. However, like an Ex-pCM, a pbMutSel model still assumes a single set of time-homogeneous site-specific amino-acid preferences for the entire tree.

Comparing ExpCM and pbMutSel models can help determine the ultimate limits of mutation-selection models that assign each site a single set of amino-acid preferences. If the limitations of ExpCMs described above arise simply because the deep mutational scanning experiments do not correctly measure the “true” amino-acid preferences across the entirety of a highly diverged phylogenetic tree, then we would expect the pbMutSel models (which infer these preferences from the entire tree) to perform better. On the other hand, if the major limitation is that no single set of time-homogenous amino-acid preferences can fully describe evolution over the entire tree, then we would expect ExpCM and pbMutSel models to perform similarly.

We fit a pbMutSel model to the entire HA phylogenetic tree, and compared the results to those from analyzing the same tree with the best ExpCM, which is the ExpCM(H1+H3 avg)+Γ*ω* variant. This is a direct apple-to-apples comparison, since the pbMutSel model also draws *ω* from a gamma-distribution (Rodrigue and Lartillot, 2014). First, we compared the amino-acid preferences inferred by the pbMutSel model to the preferences measured in the experiments. Figure 7A shows that the preferences inferred by pbMutSel are quite similar to the (H1+ H3 avg) obtained by averaging the deep mutational scanning measurements for the H1 and H3 HAs. Notably, the amino-acid preferences from the pbMutSel model are more correlated with the (H1+ H3 avg) than the H1 and H3 measurements are with each other (Figure 7A). This strong correlation indicates that the ExpCM(H1+H3 avg)+Γ*ω* is unlikely to be much different than a pbMutSel model that is parameterized only using the natural sequence data.

**Figure 7:**
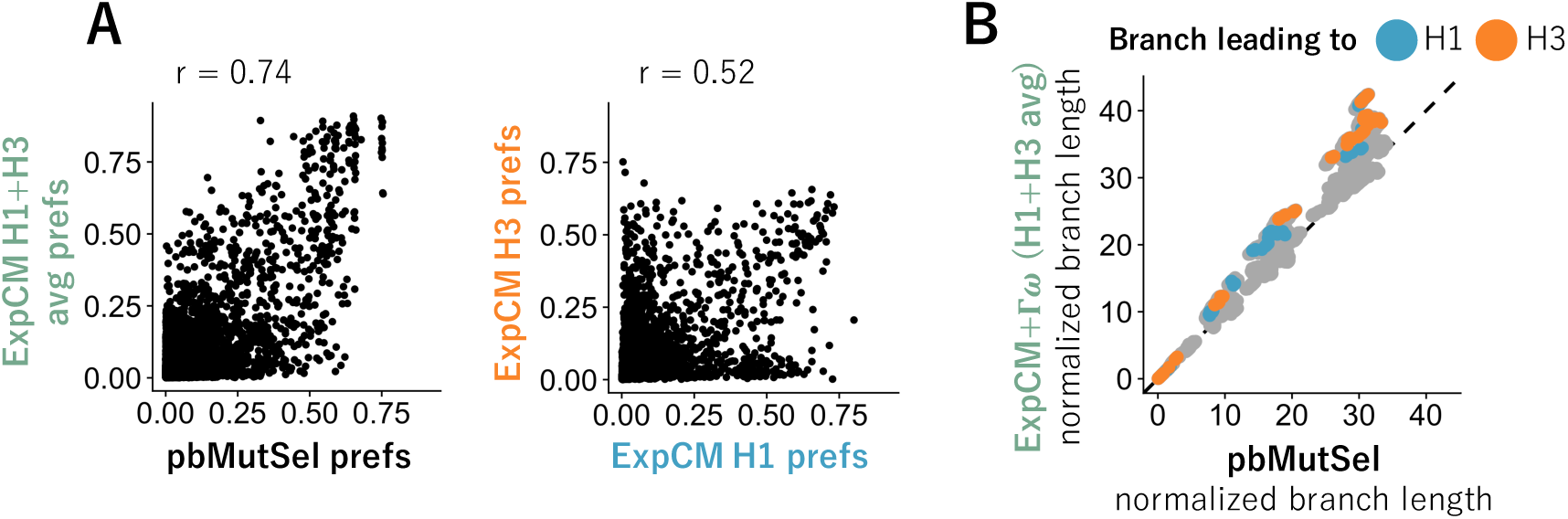
Models inferred from natural sequences have similar stationary states to models defined by experimental preferences and estimate similar branch lengths. We fit an Ex-pCM(Hl+H3 avg)+Γ*ω* and a pbMutSel to the full HA tree in Figure 4. The pbMutSel amino-acid preferences are inferred from the natural HA sequences, while the ExpCM amino-acid preferences are experimentally measured and then rescaled by the stringency parameter in Table 1. (A) The pbMutSel preferences are more correlated with the re-scaled average of the H1 and H3 deep mutational scanning preferences than the individual re-scaled H1 and H3 deep mutational scanning preferences are to each other (Pearson’s *r*: 0.74 versus 0.52). (B) The ExpCM(Hl+H3 avg)+Γ*ω* and pbMutSel models estimated similar branch lengths when fit to the entire HA tree. Points denote branch lengths between all pairs of tips on the tree. Blue and orange denote branches that lead to the H1 and H3 deep mutational scanning reference sequences respectively. The phydms program implementing ExpCMs and the `PhyloBayes–MPI` program implementing pbMutSel models give branch lengths in different units, so to facilitate direct comparison between the models, we have normalized all branch lengths returned by each program by the length of the branches separating the earliest (A/South Carolina/1918) and latest (A/Solomon Islands/2006) seasonal human H1 sequences on the tree.

We next compared the branch lengths estimated by using the ExpCM(H1+H3 avg)+Γ*ω* and pbMutSel models. As shown in Figure 7B, these two models estimated similar branch lengths across the entire HA phylogenetic tree. However, the estimates are not identical, and the tension between local and global accuracy of the amino-acid preferences is still apparent. Specifically, the branches leading to the H1 or H3 sequences used in the experiments were estimated to be slightly longer by the ExpCM, while some other branches were estimated to be slightly longer by the pb-MutSel model. The relatively longer branches leading to the experimental sequences when using the ExpCM(H1+H3 avg)+Γ*ω* suggests that the “tree average” amino-acid preferences inferred by the pbMutSel model are not as accurate as the preferences from the deep mutational scanning for sequences close to those used in the experiments. However, for sequences distant from those used in the experiments, the “tree average” preferences inferred by the pbMutSel model appear to be slightly better than the experimental values. Therefore, while the ExpCM and pbMutSel models differ slightly in the extent to which they lengthen different branches, neither model can avoid the tension between the local and global accuracy of amino-acid preferences.

## Discussion

We examined how estimates of deep branch lengths on phylogenetic trees are affected by accounting for the fact that proteins prefer specific amino acids at specific sites. We did this by comparing inferences from models informed by experimental measurements of site-specific amino-acid preferences with more conventional codon substitution models, as well as with models that infer the amino-acid preferences from the natural sequences. We found that models that account for site-specific amino-acid preferences estimated deeper long branches, regardless of whether these preferences are measured experimentally or inferred from the sequence alignment. Additionally, we showed that the extension in branch length from site-specific amino-acid preferences is mostly independent of the extension that results from simply modeling rate variation.

Overall, our results underscore the importance of modeling purifying selection in a way that is more nuanced than simply allowing the substitution rate to vary across sites. Protein sites do not simply differ in their rates of substitution—different sites also prefer different amino acids. There are now two ways to account for this fact: using models informed by deep mutational scanning experiments, or using models that infer site-specific amino-acid preferences from the natural sequence alignment. Combining either type of model with rate variation increases the inferred length of deep branches relative to models that only incorporate rate variation.

However, assuming a single set of site-specific amino-acid preferences is still an imperfect way to model evolution over a highly diverged phylogenetic tree. In the case of the experimentally informed models, it is fairly obvious why this is true: the amino-acid preferences are measured in just one genetic background, and therefore provide only a single snapshot of preferences that shift over evolutionary time due to epistasis (Pollock et al., 2012; Shah et al., 2015; Bazykin, 2015; Haddox et al., 2018; Starr et al., 2018; Doud et al., 2015). As a result, experimentally measured amino-acid preferences are most accurate for sequences similar to the one used in the experiment, and so cause the largest increases in branch length in that region of the phylogenetic tree. However, this limitation is not unique to experimentally informed models, but is a general limitation of describing purifying selection using a single set of site-independent and time-homogenous amino-acid preferences. For instance, we showed that averaging experimental measurements on two protein homologs does a somewhat better job of capturing the “average” constraint across the tree, and performs similarly to approaches that infer the “average” preferences from natural sequence data (Rodrigue et al., 2010; Rodrigue and Lartillot, 2014). But even these “average” preferences exhibit a tradeoff between local and global accuracy for the inference of deep branch lengths.

So while modeling site-specific amino-acid preferences is a clear improvement over most conventional models, the next step towards greater accuracy will require relaxing the assumption that these preferences are time homogeneous and site independent. Of course, many authors have pointed out the shortcomings of models that fail to account for the full site-interdependent complexity of purifying selection (Rodrigue et al., 2005; Choi et al., 2007; Pollock et al., 2012; Goldstein and Pollock, 2017). However, the challenge is to overcome these shortcomings with models that are tractable for real phylogenetic questions. There are two main issues: first, the Felsenstein pruning algorithm (Felsenstein, 1981) that is typically used to evaluate phylogenetic likelihoods breaks down when sites are no longer treated independently. Some alternative algorithms have been proposed (Bordner and Mittelmann, 2013; Rodrigue et al., 2009, 2005; Choi et al., 2007), but they are still in their infancy. Second, site-interdependent models require a realistic “fitness function” that describes the interactions among sites. It appears that typical structural modeling programs are insufficient for this purpose (Rodrigue et al., 2009). But hope comes from experimental progress in measuring actual site-interdependent constraints on proteins (Olson et al., 2014; Wu et al., 2016; Steinberg and Ostermeier, 2016; Li et al., 2016), combined with new methods for using these measurements to parameterize fitness functions (Sailer and Harms, 2017; Otwinowski et al., 2018; Otwinowski, 2018). Perhaps some day such truly realistic models might be useful for phylogenetic inference. Until that day, our work shows that modeling a single set of time homogenous amino-acid preferences provides at least some improvement.

## Methods

### Substitution models

All of the substitution models used in this paper have been described previously. However, here we briefly recap their exact mathematical implementations.

#### GY94 model

The GY94 model is M0 variant of the Goldman-Yang model described by Yang et al. (2000). Specifically, the substitution rate *P_xy_* from codon *x* to codon *y* is

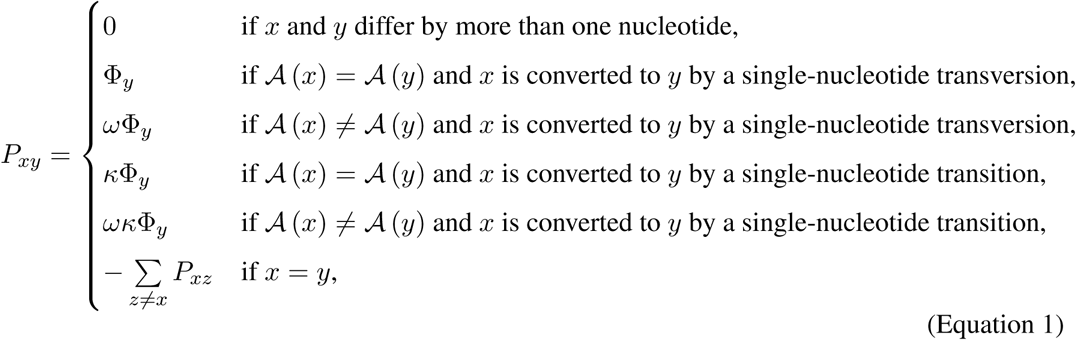

where 𝓐 (*x*) is the amino-acid encoded by codon *x*, *κ* is the transition-transversion rate, Φ_*y*_ is the equilibrium frequency of codon *y*, and *ω* is the relative rate of nonsynonymous and synonymous substitutions. We define the codon frequency parameters, Φ_*y*_, using the “corrected F3X4” method from Pond et al. (2010). There are nine parameters describing the nucleotide frequencies at each codon site (the nucleotides are constrained to sum to one at each codon position), and these parameter values are calculated from the empirical alignment frequencies. The “corrected F3X4” method calculates the Φ_*y*_ values from these nucleotide frequencies but corrects for the exclusion of sequences with premature stop codons from the analysis.

The frequency *p_x_* of codon *x* in the stationary state of a GY94 model is simply

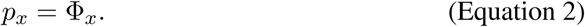

Overall, a GY94 model has 11 free parameters: *κ*, *ω*, and the 9 nucleotide frequency parameters used to define Φ_*y*_.

### Experimentally Informed Codon Model (ExpCM)

The ExpCM models used in this paper are the ones described in Bloom (2017). Briefly, the rate of substitution *P*_*r*,*xy*_ of site *r* from codon *x* to *y* is

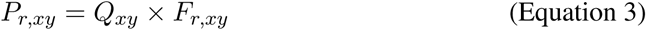

where *Q_xy_* is proportional to the rate of mutation from *x* to *y*, *F*_*r*,*xy*_ is proportional to the probability that this mutation fixes, and the diagonal elements *P_xx_* are set by *P_xx_* = −∑_*z*≠*x*_ *P_xz_*.

The rate of mutation *Q_xy_* is assumed to be uniform across sites, and takes an HKY85-like (Hasegawa et al., 1985) form as

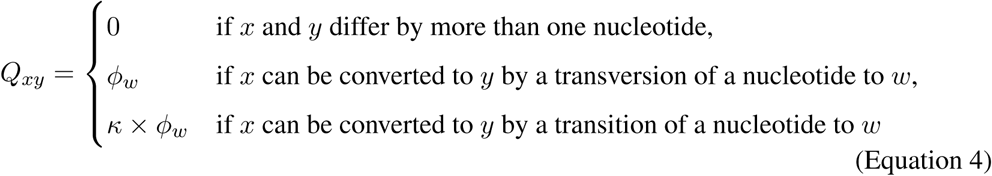

where *ϕ_w_* is the nucleotide frequency of nucleotide *ω* and *K* is the transition-transversion rate.

The deep mutational scanning amino-acid preferences are incorporated into the ExpCM via the *F*_*r*,*xy*_ terms. The experiments measure the preference *π*_*r*,*a*_ of every site *r* for every amino-acid *a*. *F*_*r*,*xy*_ is defined in terms of these experimentally measured amino-acid preferences as

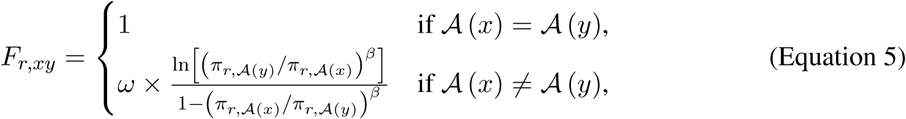

where *β* is the stringency parameter (Bloom, 2014b; Hilton et al., 2017) and *ω* is the relative rate of nonsynonymous to synonymous substitutions after accounting for the amino-acid preferences.

The stationary state of an ExpCM is

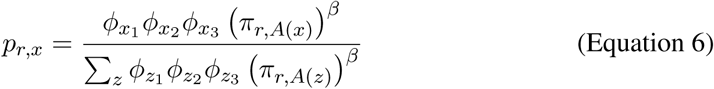

where *ϕ*_*x*_1__, *ϕ*_*x*_2__, and *ϕ*_*x*_3__ are the nucleotides at position 1, 2, and 3 of codon *x*.

An ExpCM has five free parameters: *κ*, *ω*, and the three independent *ϕ_x_* values. The amino-acid preferences *π*_*r*,*a*_ are *not* free parameters since they are determined *a priori* by an experiment independent of the sequence alignment being analyzed.

#### Γ*ω* rate variation

The GY94+Γ*ω* is equivalent to the M5 model in Yang et al. (2000) with *ω* drawn from *K* = 4 categories. The ExpCM+Γ*ω* similarly draws *ω* from a Γ distribution discretized into *K* = 4 bins. Each bin is equally weighted and *ω* takes on the mean value of the bin. Because the Γ distribution is defined by two parameters, adding Γ*ω* to a model with a single *ω* adds one free parameter. Therefore, the GY94+Γ*ω* model has 12 free parameters, and the ExpCM+Γ*ω* model has 6 free parameters.

#### GY94 with *ω_r_*

In Figure 2, we describe GY94 models where each site *r* has its own *ω_r_* value that is calculated from the amino-acid preferences using the relationship described by Spielman and Wilke (2015b). This relationship defines the expected rate of nonsynonymous to synonymous substitutions given the amino-acid preferences. We first fit an ExpCM to the “low divergence” H1 subtree (parameter values in Supplementary table 2), which allows us to calculate *P*_*r*,*xy*_ (Equation 3), *Q_xy_* (Equation 4), and *p*_*r*,*x*_ (Equation 6). We then calculated *ω_r_* using the equation of Spielman and Wilke (2015b), normalizing by the gene-wide *ω* fit by the ExpCM:

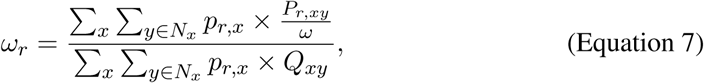

where *N_x_* is the set of codons that are nonsynonymous to codon *x* and differ from codon *x* by only one nucleotide.

### HA amino-acid preferences from deep mutational scanning experiments

We used amino-acid preferences measured in deep mutational scans of the A/WSN/1933 H1 HA (Doud and Bloom, 2016) and the A/Perth/2009 H3 HA (Lee et al., 2018) to define the amino-acid preferences that inform the ExpCMs. We only used sites that can be unambiguously aligned in these H1 and H3 HAs. These alignable sites and their mapping to sequential numbering of the HA sequences used in the deep mutational scanning experiments are in Supplementary file 1. The experimentally measured amino-acid preferences masked to just include these alignable sites are in Supplementary file 2 and Supplementary file 3. For the average preference set, we took the pairwise average of the H1 and H3 preferences. The preference for every amino acid *a* at every site *r* in the average preference set is

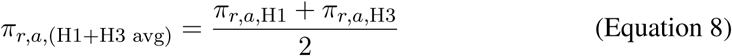

### HA sequences and tree topology

We downloaded all full-length, coding sequences for 15 of the 18 influenza A virus HA subtypes from the Influenza Virus Resource Database (Bao et al., 2008) in June of 2017. We excluded rare subtypes 15, 17, and 18, which have few sequences in the database. We filtered and aligned the sequences using `phydms_prepalignment` (Hilton et al., 2017). Specifically, we used `phydms_prepalignment` with the flag `--minidentity 0.3` to remove sequences with ambiguous nucleotides, premature stops, or frameshift mutations as well as redundant sequences. We also removed all codon sites which that are not alignable between the H1 HA and H3 HA used in the deep mutational scanning experiments (these alignable sites are listed in Supplementary file 1). We subsampled the remaining sequences to five per subtype with ≤ 1 sequence per year per subtype. We also included a small number of sequences from the major human and equine influenza lineages to ensure representation of these well-studied lineages. The resulting alignment contains 92 sequences, and is provided in Supplementary file 4.

We created four subalignments with “low” and “intermediate” divergence from either the H1 or the H3 deep mutational scanning reference sequence for the analysis in Figure 6. The “low divergence” alignments had ≥ 59% amino-acid identity to the sequence used in the deep mutational scanning, and the “intermediate divergence” alignments had ≥ 46% identity from the reference sequence (Supplementary figure 5).

We inferred the tree topology of each alignment using `RAxML` (Stamatakis, 2006) and the GTRCAT model. We estimated the branch lengths of this fixed topology using each ExpCM and GY94 models with `phydms_comprehensive` (Hilton et al., 2017).

### Asymptotic amino-acid sequence identity

For the analysis in Figure 2, we fit models to the “low divergence” H1 subtree. This gave the parameter values in Supplementary table 2.

For each model, we calculated the expected amino-acid sequence identity for two sequences separated by a branch length of *t* as

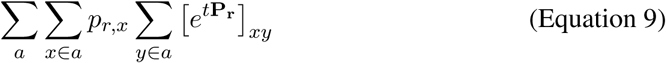

where *a* ranges over all 20 amino acids, *x* ∈ *a* indicates that *x* ranges over all codons that encode amino-acid *a*, *p*_*r*,*x*_ is the stationary state of the model at site *r* and codon *x* (given by Equation 2 for GY94-family models, and Equation 6 for ExpCM-family models), and [*e*^*t***P**_r_^]_*xy*_ is the value in row *x* and column *y* of the matrix obtained by exponentiating the product of *t* and the substitution matrix **P**_*r*_ for site *r* (defined by Equation 1 for GY94-family models and Equation 3 for ExpCM-family models).

### Simulations

For Figure 3, we simulated sequences using `pyvolve` (Spielman and Wilke, 2015a) along the full HA tree using an ExpCM defined by parameters fit to the “low divergence” H1 subtree (Supplementary table 2). We performed 10 replicate simulations and estimated the branch lengths for each replicate using `phydms_comprehensive` (Hilton et al., 2017).

### pbMutSel inference with PhyloBayes-MPI

For Figure 7, we fit a pbMutSel model to the full HA tree. We ran one chain for 5500 steps, saved every sample, and discarded the first 550 samples as a burnin. We used `PhyloBayes-MPI` program `readpb_mpi` to compute the majority-rule consensus tree and the posterior average site-specific amino-acid preferences. Convergence was assessed visually using Tracer (http://tree.bio.ed.ac.uk/software/tracer/) and by the correlation of amino-acid preferences inferred by two independent chains (r=0.996).

In order to make the branch lengths in Figure 7 comparable between the pbMutSel tree returned by `PhyloBayes-MPI` and the other trees returned by `phydms`, we we normalized the branch lengths on the pbMutSel consensus tree and the ExpCM(H1+H3 avg)+Γ*ω* by dividing each branch by the length from A/South Carolina/1/1918 and A/Solomon Islands/3/2006. These two H1 sequences are early and late representatives of the longest known human influenza lineage, and are of sufficiently high identity that different ExpCM and GY94 substitution models all estimate nearly identical branch lengths separating them.

### Software versions and computer code

All code used for the analyses in this paper is available at https://github.com/jbloomlab/divergence_timing_manuscript. External computer programs used were `phydms` (Hilton et al., 2017) version 2.2.2, `pyvolve` (Spielman and Wilke, 2015a) version 0.8.7, `PhyloBayes-MPI` (Rodrigue and Lartillot, 2014) version 1.8, `RAxML` (Stamatakis, 2006) version 8.2.11 (available at https://github.com/stamatak/standard-RAxML), `ggplot2` (Wickham, 2016), `ggtree` (Yu et al., 2017), `ggseqlogo` (Wagih, 2017), and `snakemake` (Köster and Rahmann, 2012) version 3.11.2.

## Acknowledgments

We thank Erick Matsen and Trevor Bedford for helpful comments about the project and manuscript. SKH is supported in part by training grant T32AI083203 from the NIAID of the National Institutes of Health. This work was supported by the NIAID and NIGMS of the NIH under grant numbers R01AI127893 and R01GM102198. JDB is supported in part by a Faculty Scholars grant from the Howard Hughes Medical Institute and the Simons Foundation. The funders had no role in study design, data collection and analysis, decision to publish, or preparation of the manuscript.

## Supplemental Information

**Supplementary figure 1:**
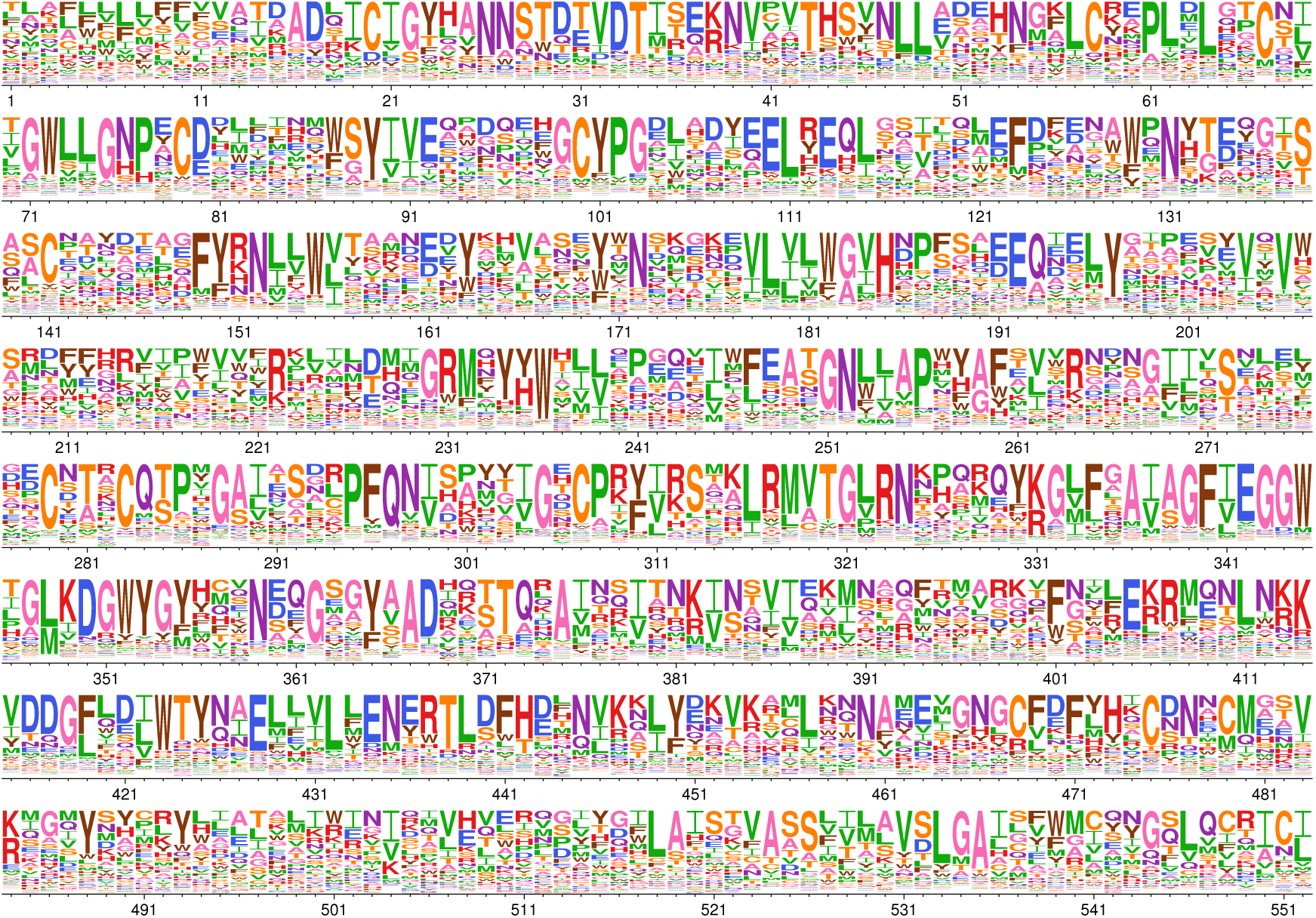
H1 HA amino-acid preferences measured by deep mutational scanning. Each column represents a site in the HA protein, and the height of each letter is proportional to the preference for the amino acid measured by Doud and Bloom (2016) and then re-scaled by the stringency parameter in Table 1. The plot only shows sites that are alignable between the H1 and H3 HAs, and these alignable sites are numbered sequentially starting from 1. The conversion between the numbering scheme in this figure and sequential numbering of the H1 HA reference sequence is in Supplementary file 1.

**Supplementary figure 2:**
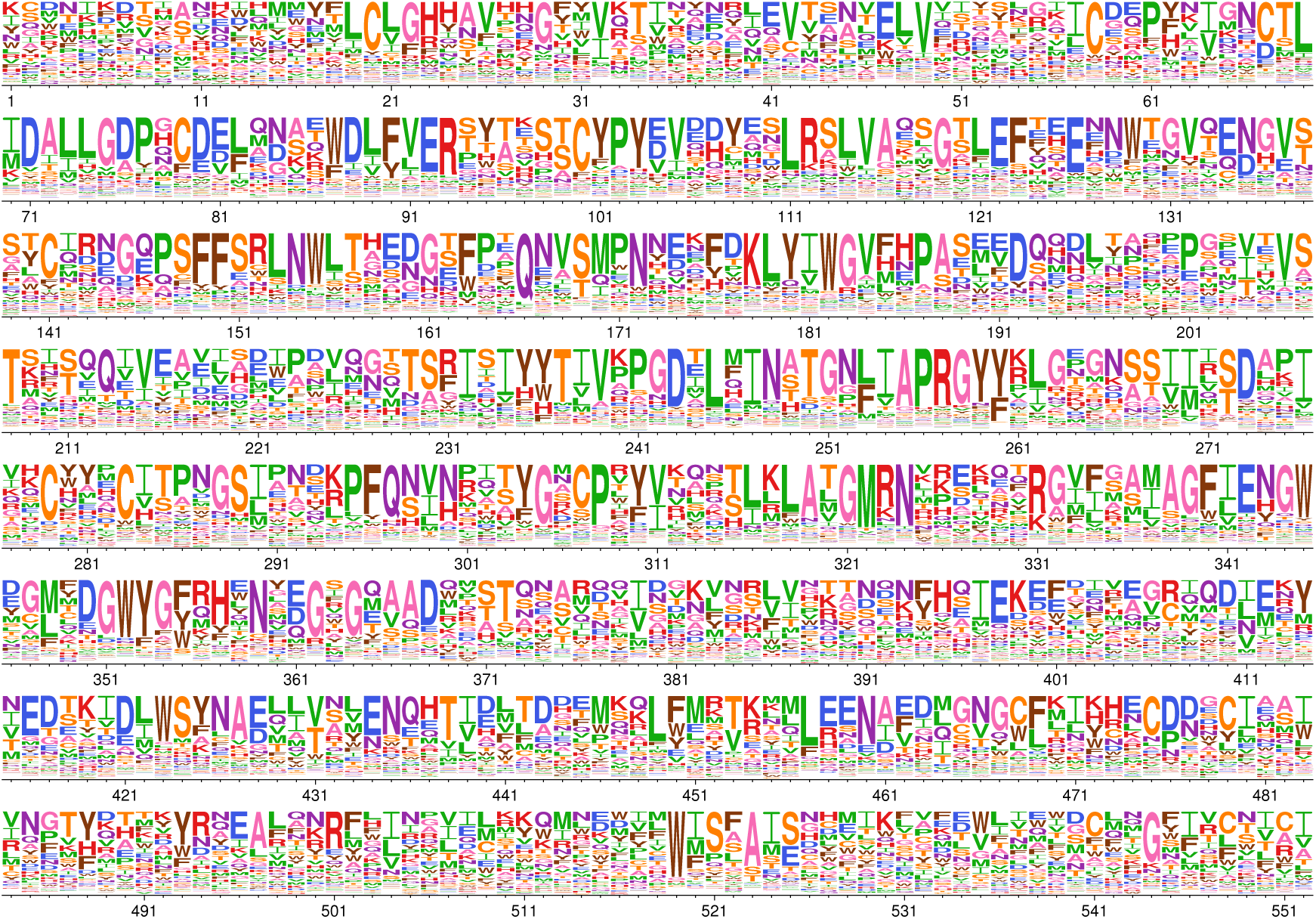
H3 HA amino-acid preferences measured by deep mutational scanning. Similar to Supplementary figure 1 but shows the re-scaled preferences for the H3 HA as measured by Lee et al. (2018).

**Supplementary figure 3:**
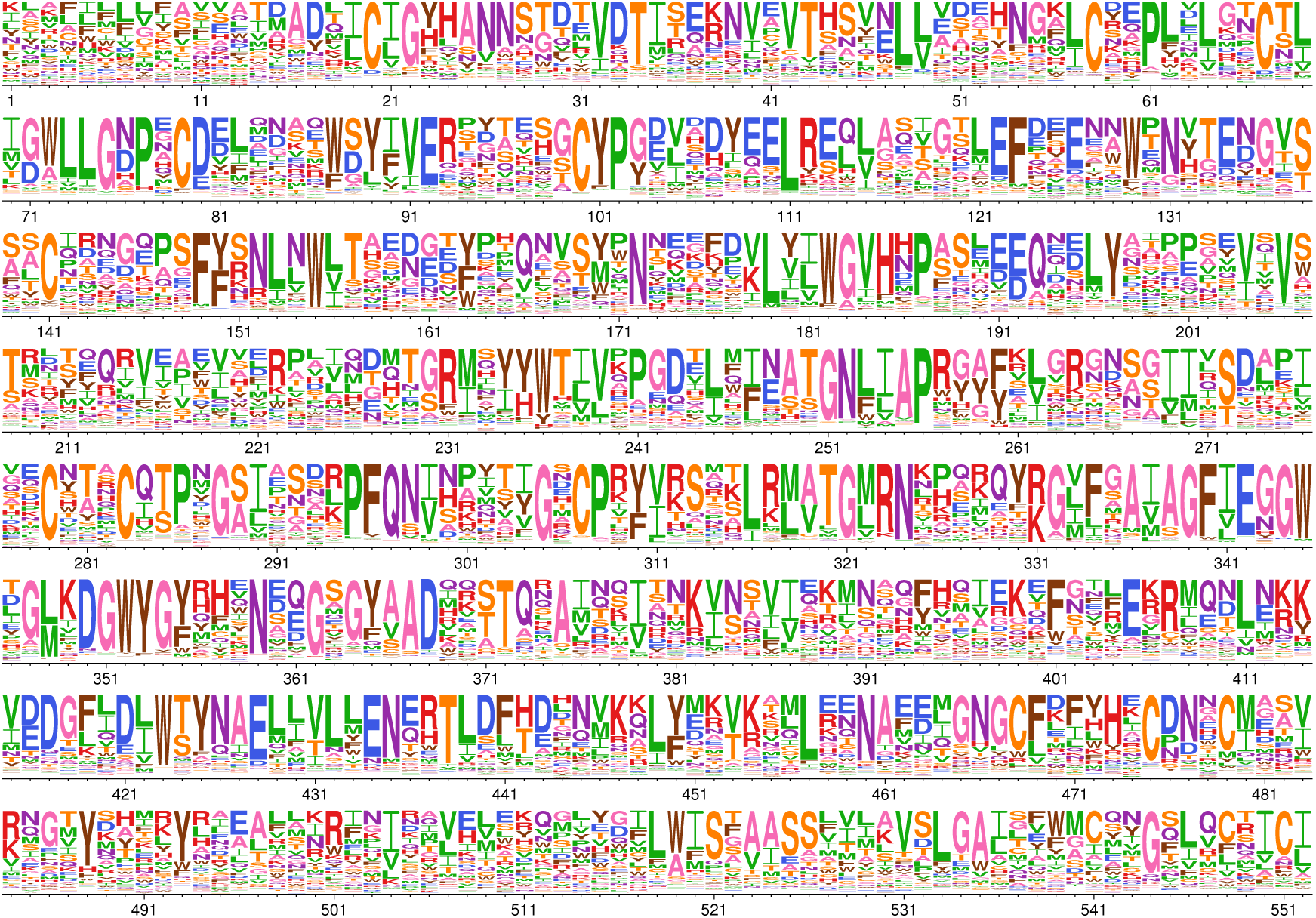
Average of H1 HA and H3 HA amino-acid preferences measured by deep mutational scanning. Similar to Supplementary figure 1 but shows the re-scaled average of the preferences for the H1 and H3 HAs.

**Supplementary figure 4:**
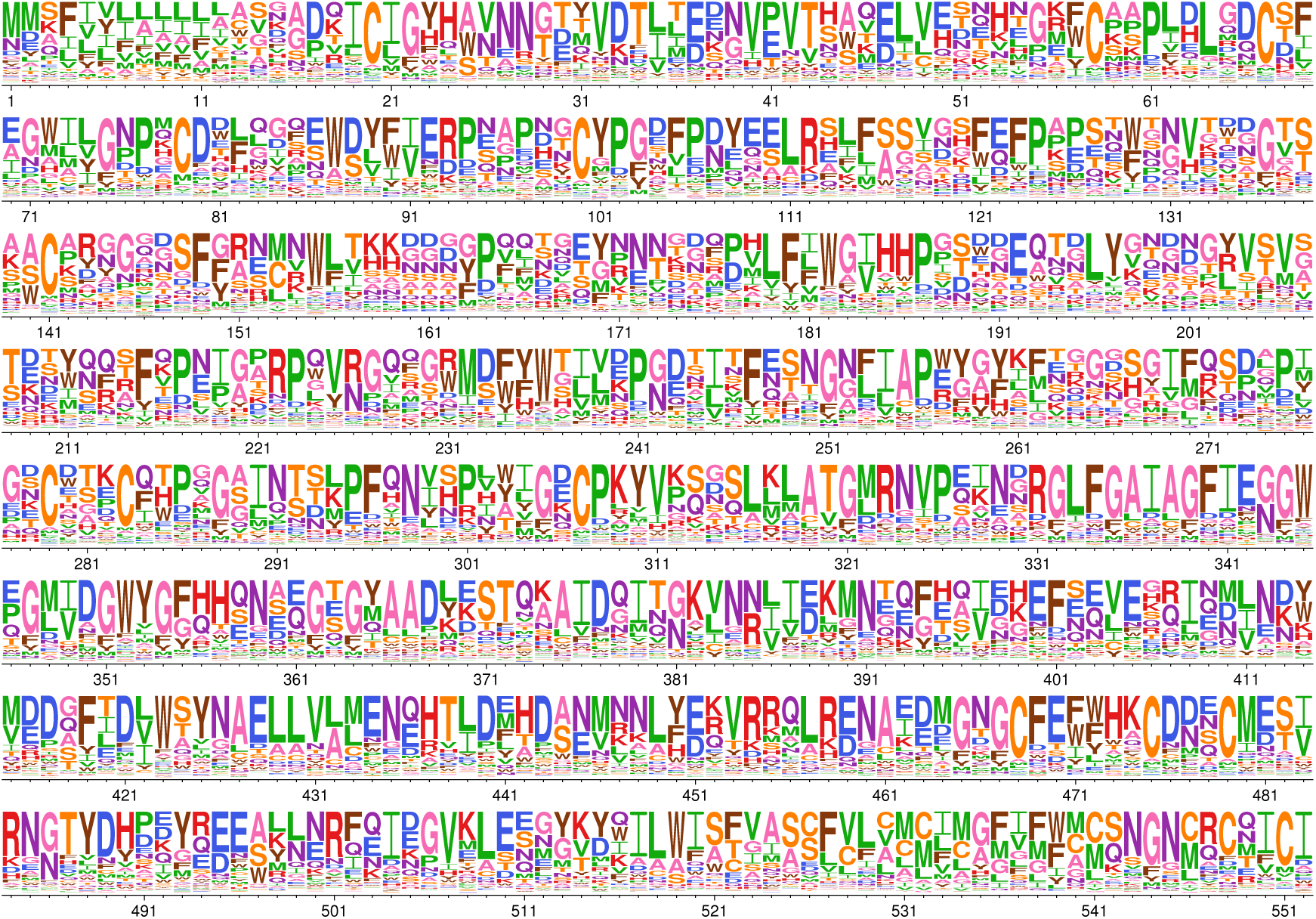
Amino-acid preferences inferred by the pbMutSel model. Similar to Supplementary figure 1, but shows the preferences inferred by fitting the pbMutSel model to the full HA tree.

**Supplementary figure 5:**
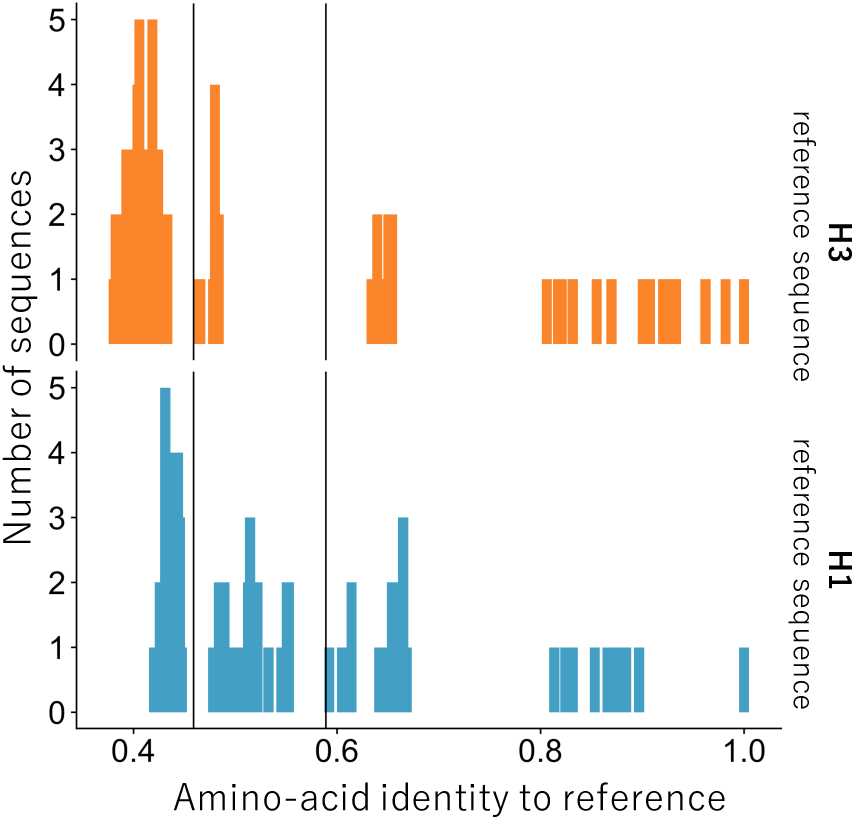
Overall divergence for subtrees. We created two subalignments for each HA used in the deep mutational scanning experiments. The “low divergence” alignments had ≥59% amino-acid identity to either the H1 or H3 reference sequence. The “intermediate divergence” alignments had ≥46% amino-acid identity to the reference sequences.

**Supplementary table 1:** Branch length extension as measured by tree diameter. We calculated the tree diameter, the distance between the two most divergent tips, for the trees in Figure 4. For each tree, the diameter is reported as a raw value and as a percentage of the GY94 model tree, the smallest of the eight trees.

| Model | Tree diameter<br>(average codon substitutions per site) | Percentage of<br>GY94 tree diameter |
| --- | --- | --- |
| GY94 | 12.04 | 100% |
| ExpCM(H1) | 14.70 | 122% |
| ExpCM(H3) | 16.28 | 135% |
| ExpCM(H1+H3 avg) | 19.21 | 160% |
| GY94 + $\Gamma\omega$ | 19.15 | 159% |
| ExpCM(H1) + $\Gamma\omega$ | 24.75 | 206% |
| ExpCM(H3) + $\Gamma\omega$ | 25.03 | 208% |
| ExpCM(H1+H3 avg) + $\Gamma\omega$ | 30.78 | 256% |

**Supplementary table 2:**
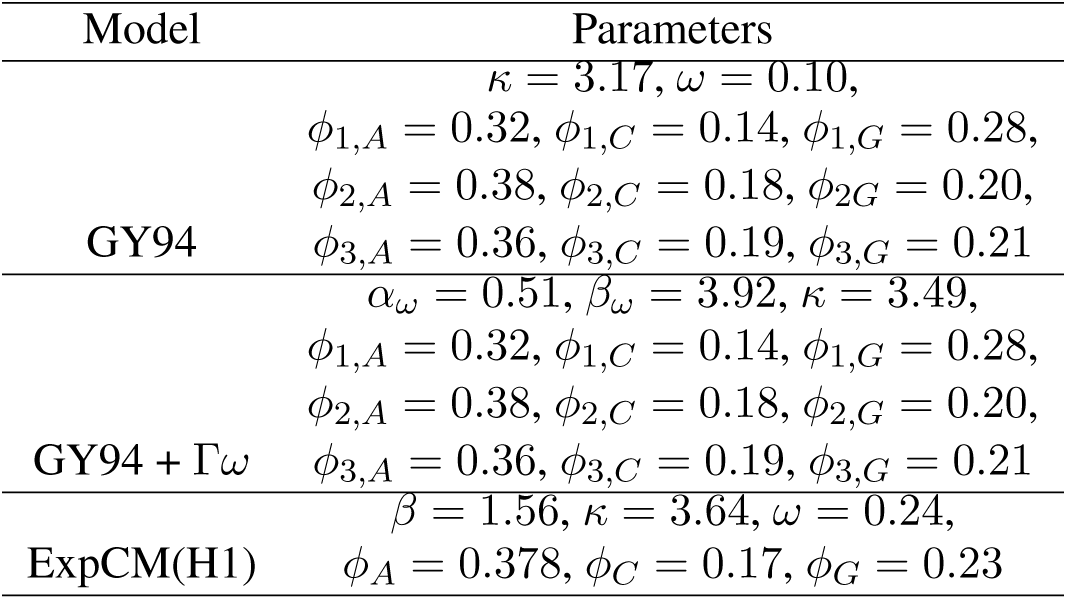
Model parameters fit to a low divergence tree. We fit GY94 models and an ExpCM defined by H1 deep mutational scanning preferences to the “low divergence from H1” tree in Figure 6. We used these model parameters calculate the expected pairwise sequence identity in Figure 2 and simulate the sequences in Figure 3.

**Supplementary file 1:** List of alignable sites between H1 HA and H3 HA. This files provides a conversion between the numbering scheme we use in the paper (sequential numbering of just the alignable sites) to sequential numbering of the H1 HA reference sequence A/Wilson Smith/1933 and the H3 HA reference sequence A/Perth/2009.

**Supplementary file 2:** Amino acid preferences measured by the deep mutational scanning of the H1 HA strain A/WSN/1933 (Doud and Bloom, 2016). This file only contains measurements for the alignable sites between H1 and H3 HAs. Conversion from this numbering scheme to sequential numbering of A/WSN/1933 is in Supplementary file 1.

**Supplementary file 3:** Amino acid preferences measured by the deep mutational scanning of the H3 HA strain A/Perth/2009 (Lee et al., 2018). This file only contains measurements for the alignable sites between H1 and H3 HAs. Conversion from this numbering scheme to sequential numbering of A/Perth/2009 is in in Supplementary file 1.

**Supplementary file 4:** The HA sequences for the full HA tree. The sequences in this alignment contain only the alignable sites between H1 and H3 HAs. Conversion from this numbering scheme to sequential numbering of A/Perth/2009 is in in Supplementary file 1.

## References

Aiewsakun P, Katzourakis A. 2016. Time-dependent rate phenomenon in viruses. Journal of Virology. 90:7184–7195.

Arenas M. 2015. Trends in substitution models of molecular evolution. Frontiers in Genetics. 6:319.

Bao Y, Bolotov P, Dernovoy D, Kiryutin B, Zaslavsky L, Tatusova T, Ostell J, Lipman D. 2008. The influenza virus resource at the National Center for Biotechnology Information. Journal of Virology. 82:596–601.

Bazykin GA. 2015. Changing preferences: deformation of single position amino acid fitness landscapes and evolution of proteins. Biology Letters. 11:20150315.

Bedford T, Suchard MA, Lemey P, Dudas G, Gregory V, Hay AJ, McCauley JW, Russell CA, Smith DJ, Rambaut A. 2014. Integrating influenza antigenic dynamics with molecular evolution. eLife. 3:e01914.

Bloom JD. 2014a. An experimentally determined evolutionary model dramatically improves phylogenetic fit. Molecular Biology and Evolution. 31:1956–1978.

Bloom JD. 2014b. An experimentally informed evolutionary model improves phylogenetic fit to divergent lactamase homologs. Mol. Biol. Evol. 31:2753–2769.

Bloom JD. 2017. Identification of positive selection in genes is greatly improved by using experimentally informed site-specific models. Biology Direct. 12:1.

Bordner AJ, Mittelmann HD. 2013. A new formulation of protein evolutionary models that account for structural constraints. Molecular Biology and Evolution. 31:736–749.

Carroll SA, Towner JS, Sealy TK, McMullan LK, Khristova ML, Burt FJ, Swanepoel R, Rollin PE, Nichol ST. 2013. Molecular evolution of viruses of the family filoviridae based on 97 whole-genome sequences. Journal of Virology. 87:2608–2616.

Choi SC, Hobolth A, Robinson DM, Kishino H, Thorne JL. 2007. Quantifying the impact of protein tertiary structure on molecular evolution. Molecular Biology and Evolution. 24:1769–1782.

Doud MB, Ashenberg O, Bloom JD. 2015. Site-specific amino acid preferences are mostly conserved in two closely related protein homologs. Mol. Biol. Evol. 32:2944–2960.

Doud MB, Bloom JD. 2016. Accurate measurement of the effects of all amino-acid mutations to influenza hemagglutinin. Viruses. 8:155.

Drummond AJ, Ho SY, Phillips MJ, Rambaut A. 2006. Relaxed phylogenetics and dating with confidence. PLoS Biology. 4:e88.

Duchêne DA, Duchêne S, Holmes EC, Ho SY. 2015a. Evaluating the adequacy of molecular clock models using posterior predictive simulations. Molecular Biology and Evolution. 32:2986–2995.

Duchêne S, Di Giallonardo F, Holmes EC. 2015b. Substitution model adequacy and assessing the reliability of estimates of virus evolutionary rates and time scales. Molecular Biology and Evolution. 33:255–267.

Duchêne S, Holmes EC, Ho SY. 2014. Analyses of evolutionary dynamics in viruses are hindered by a time-dependent bias in rate estimates. Proc. R. Soc. B. 281:20140732.

Echave J, Spielman SJ, Wilke CO. 2016. Causes of evolutionary rate variation among protein sites. Nature Reviews Genetics.

Fares MA, Holmes EC. 2002. A revised evolutionary history of hepatitis b virus (HBV). Journal of Molecular Evolution. 54:807–814.

Felsenstein J. 1981. Evolutionary trees from DNA sequences: a maximum likelihood approach. J. Mol. Evol. 17:368–376.

Fowler DM, Fields S. 2014. Deep mutational scanning: a new style of protein science. Nature Methods. 11:801–807.

Furuse Y, Suzuki A, Oshitani H. 2010. Origin of measles virus: divergence from rinderpest virus between the 11th and 12th centuries. Virology journal. 7:52.

Goldman N, Yang Z. 1994. A codon-based model of nucleotide substitution for protein-coding dna sequences. Molecular Biology and Evolution. 11:725–736.

Goldstein RA, Pollock DD. 2017. Sequence entropy of folding and the absolute rate of amino acid substitutions. Nature Ecology & Evolution. 1:1923.

Gong LI, Suchard MA, Bloom JD. 2013. Stability-mediated epistasis constrains the evolution of an influenza protein. eLife. 2:e00631.

Ha Y, Stevens DJ, Skehel JJ, Wiley DC. 2002. H5 avian and H9 swine influenza virus haemagglutinin structures: possible origin of influenza subtypes. The EMBO journal. 21:865–875.

Haddox HK, Dingens AS, Hilton SK, Overbaugh J, Bloom JD. 2018. Mapping mutational effects along the evolutionary landscape of HIV envelope. eLife. 7:e34420.

Halpern AL, Bruno WJ. 1998. Evolutionary distances for protein-coding sequences: modeling site-specific residue frequencies. Molecular Biology and Evolution. 15:910–917.

Harms MJ, Thornton JW. 2014. Historical contingency and its biophysical basis in glucocorticoid receptor evolution. Nature. 512:203–207.

Hasegawa M, Kishino H, Yano Ta. 1985. Dating of the human-ape splitting by a molecular clock of mitochondrial DNA. Journal of Molecular Evolution. 22:160–174.

Hilton SK, Doud MB, Bloom JD. 2017. phydms: Software for phylogenetic analyses informed by deep mutational scanning. PeerJ. 5:e3657.

Ho SY, Duchêne S, Molak M, Shapiro B. 2015. Time-dependent estimates of molecular evolutionary rates: evidence and causes. Molecular Ecology. 24:6007–6012.

Holmes EC. 2003. Molecular clocks and the puzzle of RNA virus origins. Journal of Virology. 77:3893–3897.

Köster J, Rahmann S. 2012. Snakemake–a scalable bioinformatics workflow engine. Bioinformatics. 28:2520–2522.

Lartillot N. 2014. The Bayesian Kitchen: overcoming the fear of over-paramerization. http://bayesiancook.blogspot.com/2014/01/the-myth-of-over-parameterization.html. Last accessed: March-12-2018.

Lartillot N, Brinkmann H, Philippe H. 2007. Suppression of long-branch attraction artefacts in the animal phylogeny using a site-heterogeneous model. BMC Evolutionary Biology. 7:S4.

Lartillot N, Philippe H. 2004. A Bayesian mixture model for across-site heterogeneities in the amino-acid replacement process. Molecular Biology and Evolution. 21:1095–1109.

Le SQ, Lartillot N, Gascuel O. 2008. Phylogenetic mixture models for proteins. Phil. Trans. R. Soc. B. 363:3965–3976.

Lee JM, Huddleston J, Doud MB, Hooper K, Wu NC, Bedford T, Bloom JD. 2018. Deep mutational scanning of hemagglutinin helps predict evolutionary fates of human H3N2 influenza variants. bioRxiv. DOI: 10.1101/298364.

Li C, Qian W, Maclean CJ, Zhang J. 2016. The fitness landscape of a trna gene. Science. 352:837–840.

McCandlish DM, Stoltzfus A. 2014. Modeling evolution using the probability of fixation: history and implications. The Quarterly Review of Biology. 89:225–252.

Murrell B, Weaver S, Smith MD, et al. (11 co-authors). 2015. Gene-wide identification of episodic selection. Molecular Biology and Evolution. 32:1365–1371.

Nielsen R. 2006. Statistical methods in molecular evolution. Springer.

Nobusawa E, Aoyama T, Kato H, Suzuki Y, Tateno Y, Nakajima K. 1991. Comparison of complete amino acid sequences and receptor-binding properties among 13 serotypes of hemagglutinins of influenza A viruses. Virology. 182:475–485.

Olson CA, Wu NC, Sun R. 2014. A comprehensive biophysical description of pairwise epistasis throughout an entire protein domain. Current Biology. 24:2643–2651.

Ortlund EA, Bridgham JT, Redinbo MR, Thornton JW. 2007. Crystal structure of an ancient protein: evolution by conformational epistasis. Science. 317:1544–1548.

Otwinowski J. 2018. Inferring protein stability and function from a high-throughput assay. arXiv. 1802.08744.

Otwinowski J, McCandlish DM, Plotkin J. 2018. Inferring the shape of global epistasis. bioRxiv. DOI: 10.1101/278630.

Philippe H, Laurent J. 1998. How good are deep phylogenetic trees? Current Opinion in Genetics & Development. 8:616–623.

Pollock DD, Thiltgen G, Goldstein RA. 2012. Amino acid coevolution induces an evolutionary stokes shift. Proc. Natl. Acad. Sci. USA. 109:E1352–E1359.

Pond SK, Delport W, Muse SV, Scheffler K. 2010. Correcting the bias of empirical frequency parameter estimators in codon models. PLoS One. 5:e11230.

Posada D, Buckley TR. 2004. Model selection and model averaging in phylogenetics: advantages of Akaike information criterion and Bayesian approaches over likelihood ratio tests. Systematic Biology. 53:793–808.

Quang SL, Gascuel O, Lartillot N. 2008. Empirical profile mixture models for phylogenetic reconstruction. Bioinformatics. 24:2317–2323.

Rodrigue N. 2013. On the statistical interpretation of site-specific variables in phylogeny-based substitution models. Genetics. 193:557–564.

Rodrigue N, Kleinman CL, Philippe H, Lartillot N. 2009. Computational methods for evaluating phylogenetic models of coding sequence evolution with dependence between codons. Mol. Biol. Evol. 26:1663–1676.

Rodrigue N, Lartillot N. 2014. Site-heterogeneous mutation-selection models within the PhyloBayes-MPI package. Bioinformatics. 30:1020–1021.

Rodrigue N, Lartillot N. 2017. Detecting adaptation in protein-coding genes using a Bayesian site-heterogeneous mutation-selection codon substitution model. Molecular Biology and Evolution. 34:204–214.

Rodrigue N, Lartillot N, Bryant D, Philippe H. 2005. Site interdependence attributed to tertiary structure in amino acid sequence evolution. Gene. 347:207–217.

Rodrigue N, Philippe H, Lartillot N. 2010. Mutation-selection models of coding sequence evolution with site-heterogeneous amino acid fitness profiles. Proceedings of the National Academy of Sciences. 107:4629–4634.

Russell R, Gamblin S, Haire L, Stevens D, Xiao B, Ha Y, Skehel J. 2004. H1 and H7 influenza haemagglutinin structures extend a structural classification of haemagglutinin subtypes. Virology. 325:287–296.

Sailer ZR, Harms MJ. 2017. Detecting high-order epistasis in nonlinear genotype-phenotype maps. Genetics. 205:1079–1088.

Shah P, McCandlish DM, Plotkin JB. 2015. Contingency and entrenchment in protein evolution under purifying selection. Proceedings of the National Academy of Sciences. 112:E3226–E3235.

Spielman SJ, Wilke CO. 2015a. Pyvolve: a flexible Python module for simulating sequences along phylogenies. PLoS One. 10:e0139047.

Spielman SJ, Wilke CO. 2015b. The relationship between dN/dS and scaled selection coefficients. Molecular Biology and Evolution. 32:1097–1108.

Stamatakis A. 2006. RAxML-VI-HPC: maximum likelihood-based phylogenetic analyses with thousands of taxa and mixed models. Bioinformatics. 22:2688–2690.

Starr TN, Flynn JM, Mishra P, Bolon DNA, Thornton JW. 2018. Pervasive contingency and entrenchment in a billion years of Hsp90 evolution. Proceedings of the National Academy of Sciences. DOI: 10.1073/pnas.1718133115.

Steinberg B, Ostermeier M. 2016. Shifting fitness and epistatic landscapes reflect trade-offs along an evolutionary pathway. Journal of Molecular Biology. 428:2730–2743.

Tamuri AU, dos Reis M, Goldstein RA. 2012. Estimating the distribution of selection coefficients from phylogenetic data using sitewise mutation-selection models. Genetics. 190:1101–1115.

Tamuri AU, Goldman N, dos Reis M. 2014. A penalized likelihood method for estimating the distribution of selection coefficients from phylogenetic data. Genetics. pp. genetics–114.

Taylor DJ, Ballinger MJ, Zhan JJ, Hanzly LE, Bruenn JA. 2014. Evidence that Ebolaviruses and Cuevaviruses have been diverging from Marburgviruses since the Miocene. PeerJ. 2:e556.

Tufts DM, Natarajan C, Revsbech IG, Projecto-Garcia J, Hoffmann FG, Weber RE, Fago A, Moriyama H, Storz JF. 2014. Epistasis constrains mutational pathways of hemoglobin adaptation in high-altitude pikas. Molecular Biology and Evolution. 32:287–298.

Wagih O. 2017. ggseqlogo: a versatile R package for drawing sequence logos. Bioinformatics. 33:3645–3647.

Wang HC, Li K, Susko E, Roger AJ. 2008. A class frequency mixture model that adjusts for site-specific amino acid frequencies and improves inference of protein phylogeny. BMC Evolutionary Biology. 8:331.

Wertheim JO, Chu DK, Peiris JS, Pond SLK, Poon LL. 2013. A case for the ancient origin of coronaviruses. Journal of Virology. 87:7039–7045.

Wertheim JO, Kosakovsky Pond SL. 2011. Purifying selection can obscure the ancient age of viral lineages. Molecular Biology and Evolution. 28:3355–3365.

Wertheim JO, Worobey M. 2009. Dating the age of the SIV lineages that gave rise to HIV-1 and HIV-2. PLoS Computational Biology. 5:e1000377.

Wickham H. 2016. ggplot2: elegant graphics for data analysis. Springer.

Worobey M, Telfer P, Souquière S, et al. (11 co-authors). 2010. Island biogeography reveals the deep history of siv. Science. 329:1487–1487.

Wu NC, Dai L, Olson CA, Lloyd-Smith JO, Sun R. 2016. Adaptation in protein fitness landscapes is facilitated by indirect paths. eLife. 5:e16965.

Yang Z. 1994. Maximum likelihood phylogenetic estimation from DNA sequences with variable rates over sites: approximate methods. J. Mol. Evol. 39:306–314.

Yang Z, Nielsen R. 2008. Mutation-selection models of codon substitution and their use to estimate selective strengths on codon usage. Molecular Biology and Evolution. 25:568–579.

Yang Z, Nielsen R, Goldman N, Pedersen AMK. 2000. Codon-substitution models for heterogeneous selection pressure at amino acid sites. Genetics. 155:431–449.

Yang Z, Rannala B. 2012. Molecular phylogenetics: principles and practice. Nature Reviews Genetics. 13:303.

Yu G, Smith DK, Zhu H, Guan Y, Lam TTY. 2017. ggtree: an R package for visualization and annotation of phylogenetic trees with their covariates and other associated data. Methods in Ecology and Evolution. 8:28–36.

Zuckerkandl E, Pauling L. 1965. Evolutionary divergence and convergence in proteins. In: Evolving genes and proteins. New York, NY: Academic Press, pp. 97–166.

